# SF3B1 promotes tumor malignancy through splicing-independent activation of HIF1α

**DOI:** 10.1101/2020.11.10.376780

**Authors:** Patrik T. Simmler, Cédric Cortijo, Lisa M. Koch, Patricia Galliker, Moritz Schäfer, Silvia Angori, Hella A. Bolck, Christina Mueller, Ana Vukolic, Peter Mirtschink, Yann Christinat, Natalie R. Davidson, Kjong-Van Lehmann, Giovanni Pellegrini, Chantal Pauli, Daniela Lenggenhager, Ilaria Guccini, Till Ringel, Christian Hirt, Gunnar Rätsch, Matthias Peter, Holger Moch, Markus Stoffel, Gerald Schwank

## Abstract

Mutations in the splicing factor SF3B1 are frequently occurring in various cancers and drive tumor progression through the activation of cryptic splice sites in multiple genes. Recent studies also demonstrate a positive correlation between expression levels of wildtype SF3B1 and tumor malignancy, but underlying mechanisms remain elusive. Here, we report that SF3B1 acts as an activator of HIF signaling through a splicing-independent mechanism. We demonstrate that SF3B1 forms a heterotrimer with HIF1α and HIF1β, facilitating binding of the HIF1 complex to hypoxia response elements (HREs) to activate target gene expression. We further validate the relevance of this mechanism for tumor progression. Monoallelic deletion of Sf3b1 impedes formation and progression of hypoxic pancreatic cancer via impaired HIF signaling, but is well tolerated in normoxic chromophobe renal cell carcinoma. Our work uncovers an essential role of SF3B1 in HIF1 signaling, providing a causal link between high SF3B1 expression and aggressiveness of solid tumors.

## INTRODUCTION

SF3B1 is a core component of the U2 small nuclear ribonucleoprotein (snRNP) complex, involved in the recognition and selection of the branchpoint sequence in RNA splicing (Wahl et al., 2009). Missense mutations in this gene are frequently found in a number of different blood cancers (e. g. myelodysplastic syndrome, chronic lymphocytic leukemia) and solid tumors (e. g. breast and pancreatic cancers, uveal melanoma) (Quesada et al., 2012; Yoshida and Ogawa, 2014). They lead to aberrant 3’ splice site usage, resulting in the creation of novel isoforms and/or nonsense mediated decay of hundreds of genes (Alsafadi et al., 2016; Darman et al., 2015; Kahles et al., 2018; Obeng et al., 2016a). For some of these genes missplicing was shown to promote tumor malignancy (i.e. *MAP3K7*, *PPP2R5A*) (Lieu et al., 2020; Liu et al., 2020; Obeng et al., 2016b), confirming the proto-oncogenic role of SF3B1. Interestingly, recent studies have also found a link between wildtype SF3B1 expression levels and tumor aggressiveness in pancreatic-, endometrial-, prostate-, liver- and breast cancer, with higher expression being associated with adverse prognosis (Alors-Perez et al., 2021; Jiménez-Vacas et al., 2019; López-Cánovas et al., 2021; Popli et al., 2020; Zhang et al., 2020). Considering that in the heart SF3B1 overexpression is induced by hypoxia (Mirtschink et al., 2015), and that solid cancers are often poorly oxygenated (Semenza, 2003), we hypothesized that upregulation of wildtype SF3B1 in tumors facilitates adaptation to hypoxia. In this study we demonstrate that SF3B1 physically interacts with hypoxia inducible factor (HIF)1α, and promotes transcriptional response to low oxygen by enabling HIF1α-HIF1β heterodimer binding to hypoxia response elements (HREs). We furthermore demonstrate the importance of this mechanism for the progression of cancers. While in pancreatic ductal adenocarcinoma (PDAC), one of the most hypoxic cancer types (Koong et al., 2000), *SF3B1* expression adversely correlates with patient survival in patients and experimental reduction of *SF3B1* levels in a mouse model severely comprises tumor development in a mouse model, in chromophobe renal cell carcinoma (chRCC), which generally grows in a normoxic environment, heterozygous loss of SF3B1 is well tolerated.

## RESULTS

### SF3B1 is a HIF target gene that positively regulates HIF pathway activity

We previously observed that SF3B1 is regulated by HIF in ischemic heart conditions (Mirtschink et al., 2015). To assess if *SF3B1* is also a HIF target gene in cancer, we first analyzed RNA sequencing (RNA-seq) data in four different solid cancers (PDAC, prostate cancer (PCA), hepatocellular carcinoma (HCC) and breast cancer (BRCA)). Supporting our assumption, we found a close correlation between the expression of *SF3B1* and the two key components of HIF signaling, *HIF1Α* and *ARNT* (*HIF1B*) in all four cancer types (Fig. 1A, B). In accordance with previous studies (Alors-Perez et al., 2021; Jiménez-Vacas et al., 2019; López-Cánovas et al., 2021; Popli et al., 2020; Zhang et al., 2020), we also observed a tendency for shorter survival in patients with higher *SF3B1* expression (Fig. 1C). Due to the well-known hypoxic environment found in PDAC (Koong et al., 2000), we then focused on this type of cancer to further investigate the role of SF3B1 in HIF signaling. In line with the hypothesis that *SF3B1* is a HIF target, we observed a strong correlation between the mean expression of 34 known HIF target genes and the expression of *SF3B*1 in human PDAC specimen (Spearman correlation coefficient = 0.52; Fig. 1D), and using a panel of patient-derived PDAC organoids and cell lines we detected a significant upregulation of SF3B1 expression upon experimental induction of hypoxia (Fig. 1E, Suppl. Fig. 1A and B). To assess if *SF3B1* is a direct target of HIF1α, we next expressed a constitutively active variant of HIF1α (HIF1α(ΔΔP)) in two cancer cell lines containing a *SF3B1*-promoter-luciferase in which the putative hypoxia-response elements (HRE) were mutated. Confirming direct regulation by HIF1α, only the wildtype *SF3B1*-promoter-luciferase reporter but not the mutant variant was activated (Fig. 1F, Suppl. Fig. 1C).

**Figure 1.**
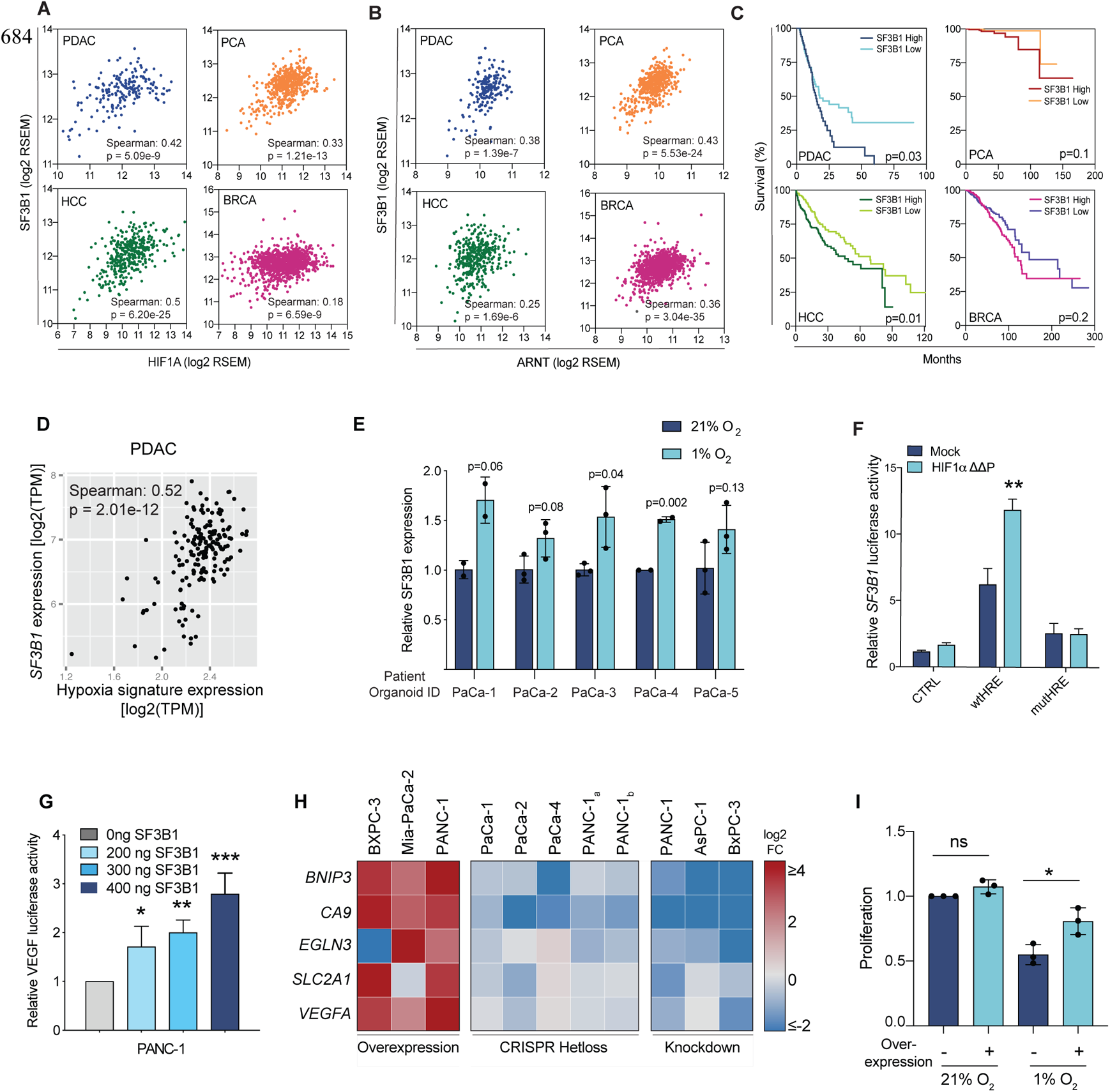
SF3B1 is required for efficient HIF signalling. **(A and B)** Scatter plot showing the correlation of *SF3B1* and *HIF1Α* (A) and *HIF1B* (B) expression in PDAC, prostate cancer (PCA), hepatocellular carcinoma (HCC) and breast cancer (BRCA). The significance of the correlation in is indicated by its p-value. (**C)** Survival of patients was stratified according to the median *SF3B1* expression in the indicated cancer types. P-values were computed with a Log-rank (Mantel-Cox) test. **(D)** Scatter plot showing the correlation between *SF3B1* mRNA levels and a hypoxia signature in PDAC patients, which is represented by the mean expression value (TPM) of 34 HIF-target genes. **(E)** SF3B1 expression in a panel of human patient-derived PDAC organoids exposed to 1% O_2_ for 12 hours. Error bars indicate standard deviation (S.D.) of biological replicates. Two-tailed unpaired t-test was used to compute the indicated p-values. **(F)** Transfection of a wildtype (WT) or HRE-mutated *SF3B1* promoter-luciferase reporter (mutHRE = AAAAA) together with either an empty vector control, or HIF1αΔΔP in PANC-1 cells. Data are shown as mean ± S.D. of biological replicates (n=3). ** P<0.01 normalized to mock wtHRE, two-tailed unpaired t-test. **(G)** Transfection of a *VEGF* promoter-luciferase reporter alone or together with either HIF1αΔΔP (the constitutively active form of HIF1α), or in combination with different concentrations of SF3B1 plasmid in PANC-1 cells. Data are shown as mean ± S.D. of biological replicates (n=3). *<P0.05, **P<0.01, ***P<0.001. One-way ANOVA followed by a Dunnet’s multiple comparison post-test. **(H)** Heatmap showing the expression of various HIF1α-target genes in patient-derived PDAC organoid lines and commercial PDAC cell lines after overexpression or knockdown of *SF3B1*, either with CRISPR-Cas9 or siRNA exposed to hypoxia for 12 hours. The indicated fold-changes are relative to the following controls: Transfection with an empty-vector (overexpression), the respective unedited cell line (CRISPR-hetloss) or treatment with control siRNA (knockdown). Expression levels are relative to the housekeeping gene *TBP*. For PANC-1, usage of two distinct guide RNAs are indicated by subscripted *a* and *b*. For each cell line and its respective control, the experiment was performed at least three times. **(I)** Impact of SF3B1 overexpression in MiaPaCa-2 cells cultured in normoxia or hypoxia for 48 hours relative to normoxic control cells (=3). *<P0.05, two-tailed unpaired t-test.

In a next step we determined whether *SF3B1* is merely a target gene or also a regulator of HIF1 signaling. We therefore manipulated *SF3B1* expression levels in multiple human PDAC cell lines and patient-derived PDAC organoids via overexpression, siRNA-mediated knock-down and monoallelic knockout (Suppl. Fig. 1D, E), and assessed its impact on HIF1 signaling. While *SF3B1* overexpression caused an increase of VEGF-luciferase reporter activity in a concentration-dependent manner (Fig. 1G, Suppl. Fig. 1F), and an upregulation of known HIF target genes (*BNIP3, CA9, EGLN3, GLUT1, VEGFA*) (Fig. 1H), reduction of *SF3B1* levels led to a significant downregulation of these target genes (Fig. 1H). In addition, the knock-down of *SF3B1* led to a 11.5-fold higher sensitivity to pharmacological inhibition of HIF1 (Suppl. Fig. 1G), and in Mia-Pa-Ca-2 cells, which are sensitive to hypoxia, overexpression of *SF3B1* increases proliferation rates in hypoxia but not in normoxia (Fig. 1I). Taken together, these data indicate that SF3B1 is not only a target gene but also a modulator of HIF1-signalling, facilitating adaptation of cancer cells to a hypoxic environment.

### SF3B1 directly interacts with HIF1*α*

SF3B1 is a well-known splice factor, and it is therefore feasible that a decrease in SF3B1 levels leads to mis-splicing of HIF pathway components. However, upon siRNA-mediated knock-down of *SF3B1* we did not observe a reduction in protein levels of the two core components of HIF signaling, HIF1α and HIF1β (Fig. 2C and 3A), and while heterozygous loss of *SF3B1* in PANC-1 cells already led to a significant reduction in HIF response, the 50% decrease in *SF3B1* levels did not lead to defective splicing in a panel of genes prone to intron retention upon *SF3B1* suppression (Paolella et al., 2017) (Suppl. Fig. 1H). Our results therefore indicate that SF3B1 may modulate HIF signaling directly in a splicing-independent manner, prompting us to assess whether SF3B1 physically interacts with HIF1α or HIF1β. Immunoprecipitation assays indeed demonstrated that under hypoxic conditions SF3B1 binds to the HIF1α/HIF1β transcriptional complex (Fig. 2A, B). While depletion of SF3B1 by specific siRNAs did not compromise binding of HIF1β to HIF1α (Fig. 2C, Suppl. Fig. 1I), siRNA-mediated depletion of HIF1α prevented co-immunoprecipitation of SF3B1 with HIF1β (Fig. 2D, Suppl. Fig. 1J). These data suggest that SF3B1 binds the HIF1α-HIF1β heterodimer via HIF1α, but that this binding is not critical for HIF1α-HIF1β heterodimer formation. To validate these findings, we produced recombinant SF3B1, HIF1α and HIF1β *in vitro* in Sf9 insect cells and mixed the purified proteins prior to analyzing their interactions *in vitro*. In this setting, SF3B1 co-immunoprecipitated with HIF1α and vice versa, irrespective whether HIF1β was present or not (Fig. 2E, F), but did not co-immunoprecipitate with HIF1β (Fig. 2F). Next, we aimed to identify the relevant domain in HIF1α that mediates binding to SF3B1 by producing a series of deletion mutants of HA-tagged HIF1α and by co-immunoprecipitating them with endogenous SF3B1 (Fig. 2G). Our results revealed that the bHLH-PAS-A-PAS-B domain is necessary and sufficient for complex formation with SF3B1 (Fig. 2H).

**Figure 2.**
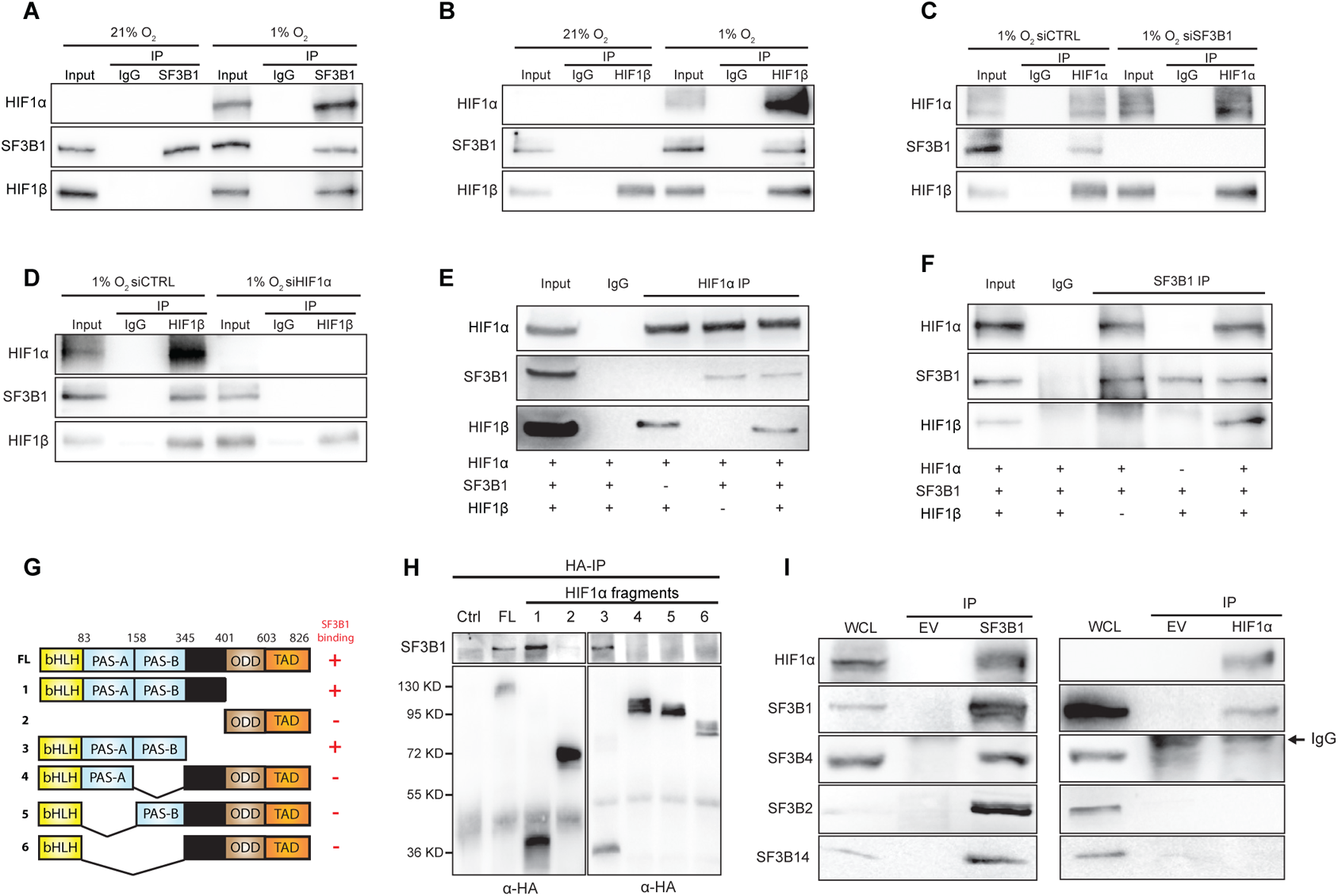
SF3B1 physically interacts with HIF1α. **(A)** Protein extracts from HEK293T cells exposed to 21% or 1% O_2_ for 12 hours were subjected to immunoprecipitation (IP) with an anti-SF3B1 antibody or anti-mouse IgG and processed for immunoblotting for indicated proteins. **(B)** Protein extracts from HEK293T cells exposed to 21% or 1% O_2_ for 12 hours were subjected to IP with an anti-HIF1β antibody or anti-mouse IgG and processed for immunoblotting for indicated proteins. **(C)** Protein extracts from HEK293T cells transfected with the indicated siRNAs, exposed to 1% O_2_ for 12 hours, were subjected to IP with an anti-HIF1α antibody or anti-rabbit IgG and processed for immunoblotting for indicated proteins. **(D)** Protein extracts from HEK293T cells transfected with the indicated siRNAs, exposed to 1% O_2_ for 12 hours, were subjected to IP with an anti-HIF1β antibody or anti-rabbit IgG and processed for immunoblotting for indicated proteins. **(E)** Recombinant strep-tagged HIF1α, SF3B1 and HIF1β proteins purified from Sf9 insect cells were mixed and subjected to IP using an anti-HIF1α antibody or anti-rabbit IgG as control and processed for immunoblotting for indicated proteins. (+) means that the indicated recombinant protein was added to the IP, (-) means that the indicated recombinant protein was omitted from the IP. **(F)** Recombinant strep-tagged HIF1α, SF3B1 and HIF1β proteins were subjected to IP using an anti-SF3B1 antibody or anti-mouse IgG as control and processed for immunoblotting for indicated proteins. (+) means that the recombinant protein was added to the IP, (-) means that the recombinant protein was omitted from the IP. **(G)** Schematic of full-length HIF1α and corresponding deletion mutants (numbered 1-6) used in IP experiments with an anti-SF3B1 antibody. (+) and (-) signs denote SF3B1 binding (experimental data are displayed in Fig. 2H). bHLH: basic helix-loop-helix, PAS-A and PAS-B: Per-ARNT-Sim domains, ODD: oxygen-dependent degradation domain and TAD: transactivation domain. **(H)** HEK293T cells were transfected with a HA-tagged WT full length (FL) or mutant forms of HIF1α (1-6). Lysates were then processed for immunoprecipitation with an anti-HA antibody (HA-IP) followed by immunoblotting with an anti-HA antibody. **(I)** Protein extracts from HEK293T cells exposed to 1% O_2_ for 8 hours were subjected to IP, with an anti-SF3B1 antibody or an anti-mouse IgG (left), or with an anti-HIF1α antibody and an anti-rabbit IgG (right). All samples were processed for immunoblotting for indicated proteins. The arrow on the right indicates the IgG band.

**Figure 3.**
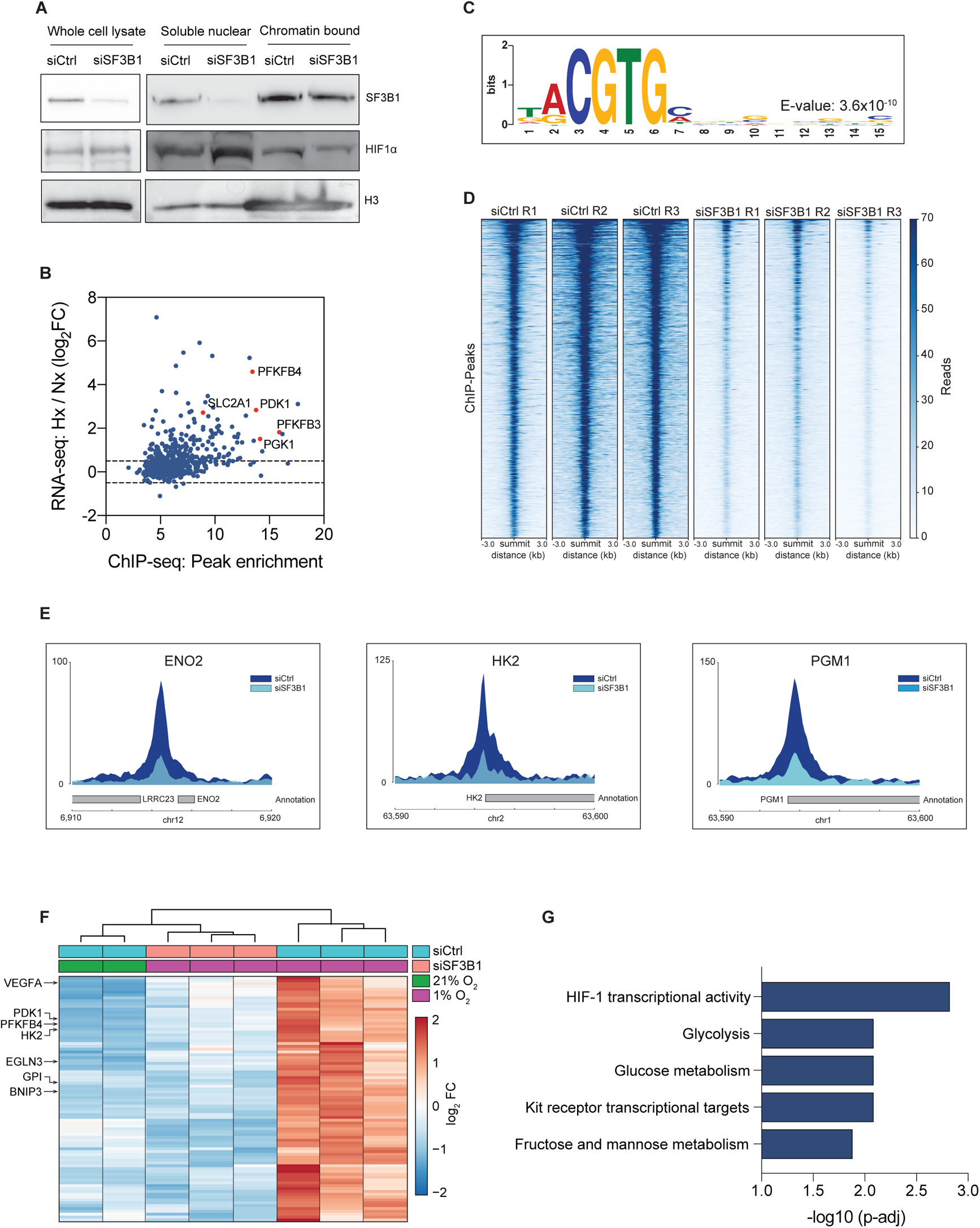
SF3B1 is essential for binding of HIF1α to its target genes. **(A)** Subcellular protein fractionation experiment in PANC-1 cells treated with control siRNA (siCtrl) or an siRNA targeting *SF3B1* (siSF3B1) and incubated in 1% O_2_ for 8 hours. SF3B1, HIF1α and H3 isolated from the indicated fractions are immunoblotted. (**B**) Enrichment of peaks identified by HIF1α-ChIP-seq plotted against the increase in expression levels (normoxia vs 1% O_2_ for 8 hours) of the corresponding nearest genes as measured by RNA-seq. **(C)** Identification of the HRE-motif (RCGTG) by MEME motif-analysis of the ChIP-seq peaks in siCtrl-treated PANC-1 cells. Nucleotide sequences of 100 bases surrounding the peak summits were analysed, representative result of one replicate is shown. **(D)** HIF1α-ChIP-seq reads of all identified peaks (n=740) of PANC-1 cells treated either with siCtrl or siSF3B1. **(E)** Peak-enrichment of the transcription start site of ENO2, HK2 and PGM2 in PANC-1 cells treated either with siCtrl or siSF3B1. Representative data from corresponding experimental replicates is shown. **(F)** Heatmap depicting nearest genes of HIF1α-ChIP-seq peaks that are upregulated in hypoxia (FC>0.5, FDR<0.1) and downregulated in siSF3B1 treated PANC-1 cells (FC<-0.5, FDR<0.1). **(G)** Pathway analysis of the gene-set shown in (F) based on the BioPlanet database. Pathways are ranked according to the adjusted P-value (p-adj).

Direct binding to the transcription factor HIF1α supports our assumption that SF3B1 modulates HIF signaling independent from its role in the spliceosome complex. To further investigate this hypothesis, we immunoprecipitated SF3B1 and HIF1α from hypoxic HEK293 cells, and immunoblotted for the presence of core components of the U2 snRNAP complex. Indeed, only SF3B1, but not HIF1α, immunoprecipitated with the U2 components SF3B2, SF3B4 and SF3B14 (Fig. 2I). Confirming these findings, HA-tagged HIF1α in PANC-1 cells co-immunoprecipitated with SF3B1, but not with SF3B2, SF3B4 and SF3B14 (Suppl. Fig. 1K).

Notably, we also assessed whether frequently occurring oncogenic mutations in SF3B1 could lead to enhanced binding with HIF1α, thereby improving the ability of cancer cells to adapt to hypoxia. However, when HIF1α was co-immunoprecipitated with oncogenic SF3B1^K700E^, we observed slightly decreased binding compared to wildtype SF3B1, and reduced HIF target gene expression under hypoxia (Suppl. Fig. 2 A-D). Furthermore, gene expression analysis in patients with solid cancers showed a consistent trend for lower expression of HIF-target genes in tumors containing *SF3B1^K700E^* (Suppl. Fig. 2E).

### DNA-binding of HIF1 to hypoxia response elements requires SF3B1

Subcellular fractionation analysis revealed that siRNA-mediated knock-down of SF3B1 leads to reduced chromatin-bound HIF1α, while leaving the total amount of HIF1α unaffected (Fig. 3A). This finding prompted us to speculate that the interaction between HIF1α and SF3B1 is essential for the binding of the HIF1 complex to its DNA target sites. Indeed, when we performed chromatin immunoprecipitations (ChIP) experiments we found that SF3B1 is bound to HREs of various HIF target genes under hypoxic conditions (Suppl. Fig. 3A), and that this binding is abolished upon knockdown of HIF1α (Suppl. Fig. 3B). Likewise, hypoxia-dependent binding of HIF1α and HIF1β to HREs was strongly reduced on upon siRNA-mediated *SF3B1* depletion (Suppl. Fig. 3C – F). To further assess SF3B1-dependency for HIF1 DNA binding in more depth, we performed HIF1α-ChIP-sequencing on hypoxic PANC-1 cells treated with an SF3B1-targeting siRNA or a control siRNA. We identified 740 high-confidence ChIP-seq peaks that were present in all three control replicates (Suppl. Table 1). Importantly, a major fraction of these peaks was identified within close proximity (± 3kb) of genes that were transcriptionally upregulated in hypoxia in PANC-1 cells under hypoxia (Fig. 3B), with 6 of the 10 most pronounced peaks being located at the transcriptional start site of the well-known HIF1 targets ENO1, BNIP3L, PFKFB3, PGAM1, PGK1 and PDK1 (Suppl. Fig. 3G). In addition, *de novo* motif analysis in the regions ± 50 bp from the peak-summits identified the previously-described RCGTG HRE-consensus sequence (Greenald et al., 2015) (Fig. 3C). In line with our hypothesis that SF3B1 is required for HIF1α binding to its DNA target sites, we observed a strong reduction in the HIF1α ChIP-seq peak-height upon *SF3B1* knock-down (Fig. 3D, E), with 99,73% of the 740 identified peaks being significantly reduced (FDR < 0.05, FC < −1.5) (Suppl. Fig. 3H, Suppl. Table 1). Differential RNA-seq in PANC1 cells further enabled us to define a subset of direct HIF1α target genes (with HIF1α-ChIP-peaks within a 3kb distance) that are dependent on SF3B1 for transcriptional upregulation upon hypoxia (Fig. 3F). As expected, unbiased pathway analysis revealed an enrichment of genes associated with HIF1 transcriptional activity and glucose metabolism (Fig. 3G), demonstrating the importance of SF3B1 for metabolic adaptation to hypoxia.

### Heterozygosity of *Sf3b1* restricts PDAC growth through impaired HIF-signalling

To investigate the importance of SF3B1-mediated HIF regulation for PDAC, which is known for its hypoxic environment and the necessity to upregulate glycolysis for tumor progression, we engineered a conditional *Sf3b1* knockout mouse model where we reduced SF3B1 levels by heterozygous deletion of *Sf3b1* (Fig. 4A). The allele was crossed into KPC (*LSL-Kras^G12D/+^; LSL-Trp53^R172H/+^; Ptf1a-Cre*) mice, where pancreas-specific activation of oncogenic *Kras^G12D^* and dysfunctional *Trp53* drives PDAC development within 9-14 weeks (Hingorani et al., 2005). As expected, homozygous deletion of *Sf3b1* in the pancreas was not viable, and heterozygous deletion in the KPC background (KPCS) led to a significant reduction in *Sf3b1* gene expression and protein levels (Fig. 4B, Suppl. Fig. 4A, B). While no abnormalities with respect to anatomical appearance and weight of the pancreas were observed in *Sf3b1^+/-^* mice in the non-cancerogenic *Ptf1a-Cre* background (Suppl. Fig. 4C, D), in the KPC background heterozygous deletion of *Sf3b1* led to significant morphological differences (Suppl. Fig. 4E). At 7 weeks of age 60% of KPC mice but only 2% of the KPCS mice developed PanIN lesions (Suppl. Fig. 2F-I), and at 13 weeks of age 100% of KPC mice developed PanIN lesions or PDAC whereas only 60% of KPCS mice had grade 1 or 2 PanIN lesions (Fig. 4C, D; Suppl. Fig. 4G). Furthermore, KPC mice developed larger tumors (Suppl. Fig. 4J, K), and were associated with considerably shorter survival (Fig. 4F).

**Figure 4.**
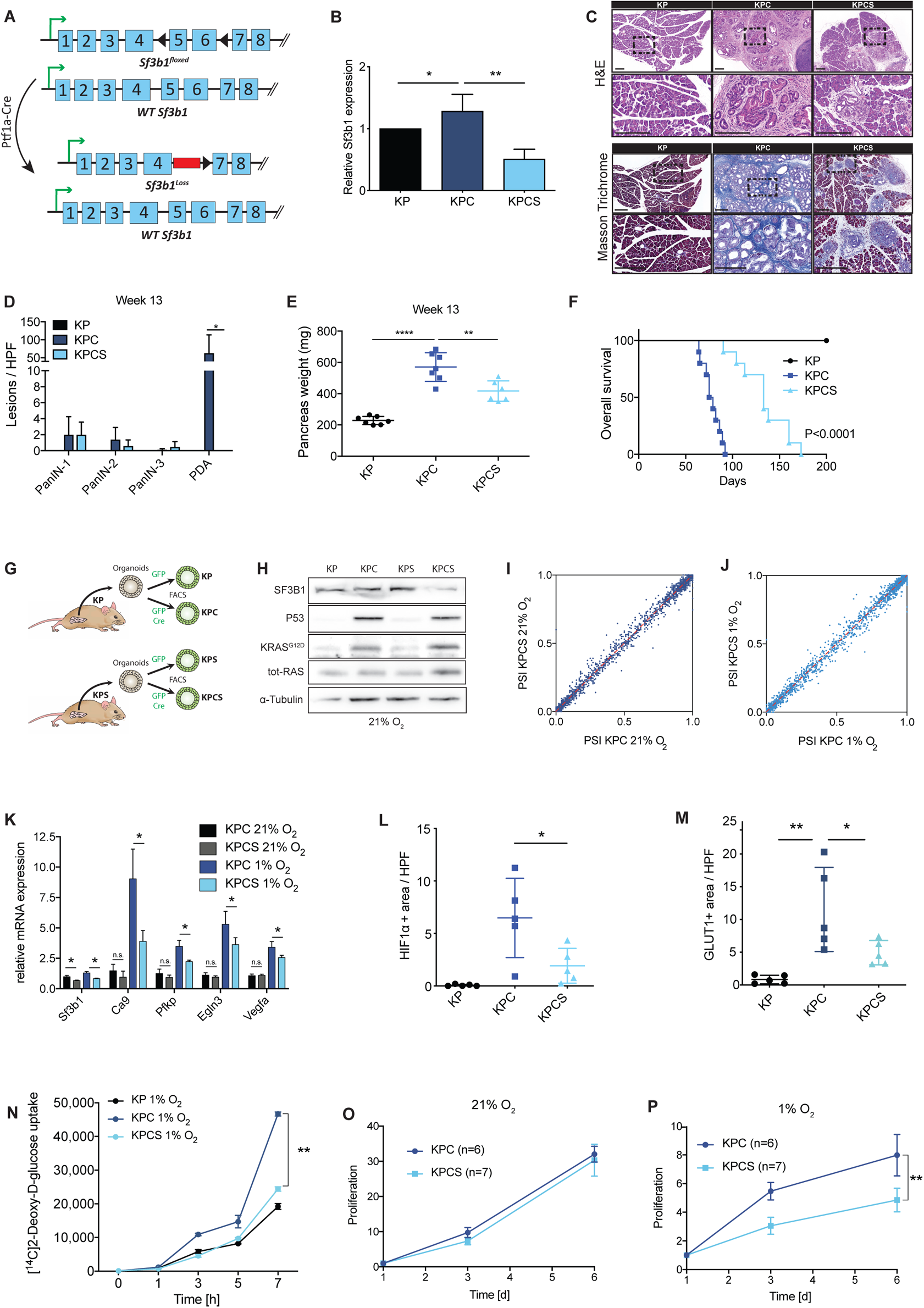
Heterozygosity of *Sf3b1* hampers PDAC progression through impaired HIF-signalling. **(A)** Schematic representation of the transgenic approach to obtain pancreas-specific heterozygosity of Sf3b1. **(B)** qPCR analysis of *Sf3b1* mRNAs from pancreata of KP, KPC and KPCS mice of 13 weeks of age. For *P<0.05 **P<0.01. One-way ANOVA followed by a Tukey’s multiple comparison post-test. **(C)** Representative images of H&E staining and Masson’s Trichrome staining showing histopathologic lesions in pancreata of KP, KPC and KPCS mice at 13 weeks of age. Dashed lines indicate the area magnified below. Scale bar is 100 µm. **(D)** Quantification of the prevalence of mPanIN and PDAC in the H&E-stained pancreatic sections of 13-week-old KP, KPC and KPCS mice. Ten high power fields (HPF) were analysed per animal. Data are represented as mean ± S.D. (n = 5). * P<0.05, ** P<0.01, *** P<0.001. One-way ANOVA followed by a Tukey’s multiple comparison post-test. **(E)** Pancreas weight of 13 weeks old KP, KPC and KPCS mice. The data are represented as mean ± S.D. (n = 6-7). ** P<0.01, **** P<0.0001. One-way ANOVA followed by a Tukey’s multiple comparison post-test. **(F)** Kaplan-Meier survival curve of KP, KPC and KPCS mice over time (KP n = 8, KPC n = 10, KPCS n = 10). Long rank test. **(G)** Schematic representation of the generation of murine tumor organoids with GFP or GFP-Cre viruses derived from KP and KPS mice. Expression of Cre recombinase in KPC and KPCS organoids leads to the activation of *Kras^G12D^* and *Trp53^R17H^* and to a loss of one allele of *Sf3b1* in KPCS organoids. **(H)** Protein extracts from KP, KPC, KPS, and KPCS organoids grown under normoxic conditions were immunoblotted for the indicated proteins. **(I and J)** Scatter plot of percent spliced-in (PSI) events in KPC and KPCS organoids cultured in normoxia (G) and hypoxia (H), representing a measure of high confidence alternative splice events (n ≥ 4 organoid lines, lines established from individual mice). **(K)** Expression levels of HIF1α target genes determined by qPCR of KPC and KPCS organoids (n = 6 organoid lines) cultured in normoxia and hypoxia for 48 hours. Expression levels are relative to the housekeeping gene *Actb*. Data are shown as mean ± SEM. For each organoid line, the experiment was performed in three biological replicates. * P<0.05, One-way ANOVA followed by a Tukey’s multiple comparison post-test. **(L and M),** Quantification of HIF1α (L) or GLUT1 (M) positive areas in KP, KPC and KPCS mice at 13 weeks of age. Ten high power fields (HPF) were analysed per animal (n = 5). * P<0.05. One-way ANOVA followed by a Tukey’s multiple comparison post-test. (**N**) KPC and KPCS organoids were incubated in medium containing [^14^C] 2-deoxyglucose at different time-points and processed for glucose uptake measurements. Organoids were exposed to 1% O_2_ for 12 hours. Counts were normalized to the cell number (*n* = 4 biological replicates per time point and condition). ** P<0.01, One-way ANOVA followed by a Tukey’s multiple comparison post-test. **(O and P)** Proliferation of organoids in normoxia (O) and hypoxia (P). Data are shown as mean ± SEM of organoid lines (n ≥ 6 organoid lines). For each organoid line, the experiment was performed in three biological replicates. ** P<0.01, indicating comparison of KPC and KPCS, Two-way ANOVA followed by a Sidak’s multiple comparisons test.

While we hypothesized that heterozygous *Sf3b1* deletion inhibits PDAC progression via attenuating HIF activity, impaired splicing could provide an alternative explanation. To discriminate between both scenarios, we established 3D organoid cultures from KP- and KPS animals (Fig. 4G, Suppl. Fig. 5A). This enabled us to analyze ductal tumor cells without stromal cell contamination (Boj et al., 2015), and to control the oxygen concentration of the environment. We first confirmed that lentiviral transduction of a CRE recombinase induced expression of KRAS^G12D^, homozygous deletion of *p53*, and heterozygous deletion of *Sf3b1,* leading to a reduction in *Sf3b1* mRNA and protein levels (Fig. 4H; Suppl. Fig 5B-D). We next analyzed global splicing efficiencies in KPC and KPCS organoids under normoxic and hypoxic conditions, but did not find substantial differences in the percentage of ‘spliced-in’ (PSI) events, including exon skips, alternative 3’and 5’ sites, and intron retentions (Fig. 4I, J; Suppl. Fig. 5E-L). In contrast, when we assessed the HIF pathway activity, we observed reduced expression of HIF target genes upon heterozygous *Sf3b1* loss under hypoxia, but not under normoxia (Fig. 4K; Suppl. Fig. 5M). These findings were confirmed *in vivo* by immunohistochemistry in mouse tumors, where the expression of HIF1α as well as the HIF target GLUT1 was reduced in KPCS mice compared to KPC mice (Fig. 4L, M; Suppl. Fig. 5N-Q), and in a functional assay for glycolysis. Incubation of organoids with radiolabeled [^14^C]-2-Deoxy-D-glucose showed no major differences in glucose uptake between KPC and KPCS organoids under normoxic conditions (Suppl. Fig. 3R), but a decrease in glucose uptake in KPCS organoids under hypoxia (Fig. 4N). Supporting these results, KPCS organoids grew significantly slower than KPC organoids in hypoxia but not in normoxia (Fig. 4O, P). Together, our results demonstrate that heterozygous deletion of *Sf3b1* in PDAC does not lead to dysfunctional splicing, but hampers adaptation to hypoxia, highlighting the importance of the SF3B1-HIF1α axis in pancreatic cancer pathogenesis.

**Figure 5.**
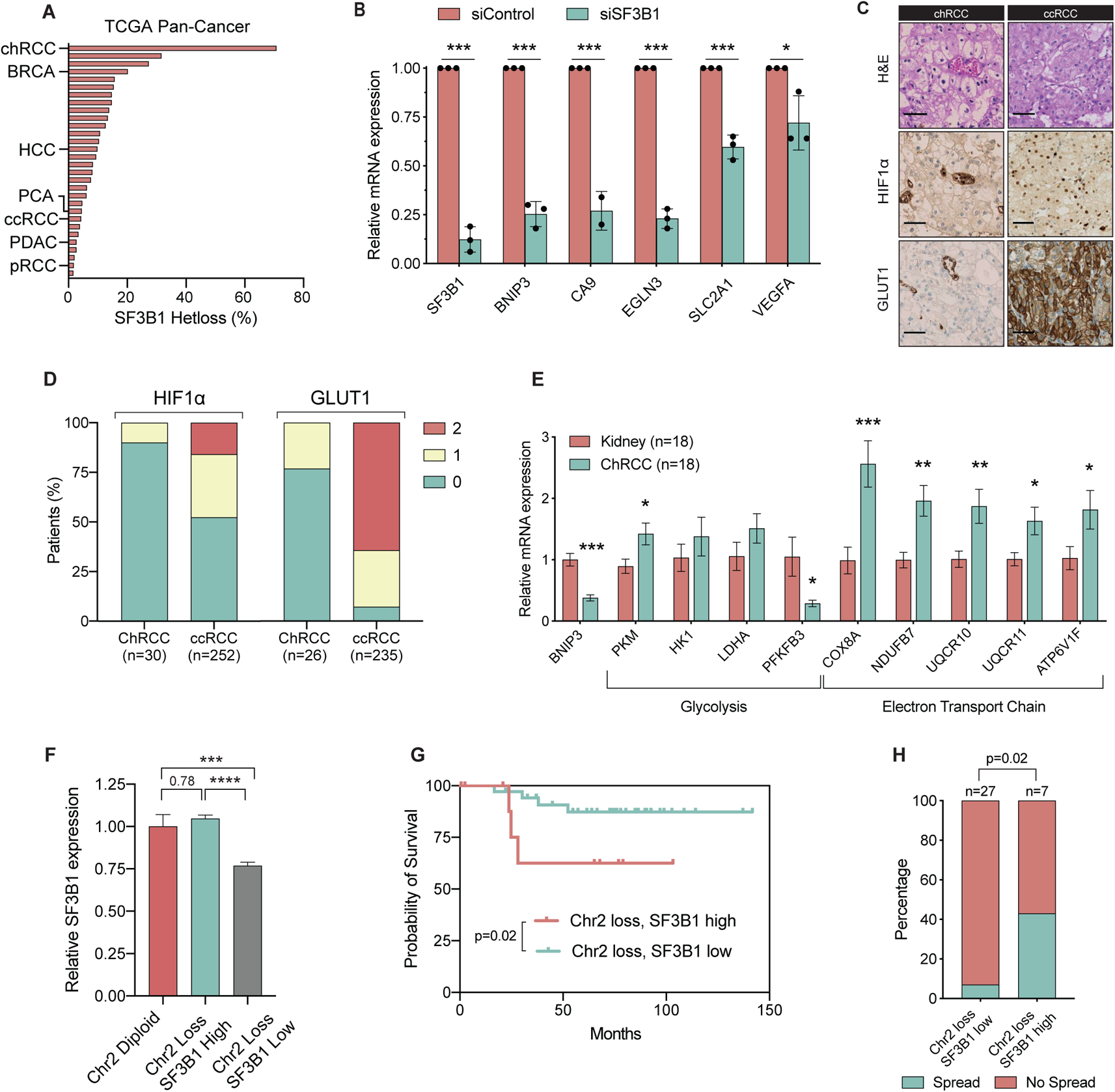
ChRCC with heterozygous SF3B1 loss grows in a normoxic environment. **(A)** Loss of heterozygosity (LOH) of SF3B1 in 30 solid tumors of the TCGA Pan-Cancer Atlas. Highlighted are the tumor types analysed in Fig 1 A-C, as well as chRCC and ccRCC. The full list is provided in Suppl. Table 2 **(B)** qPCR analysis of indicated genes in patient-derived cell chRCC cells treated with siControl or siSF3B1 under 1% O_2_ for 8 hours. Data represents results of three independent experiments. Expression levels are relative to the housekeeping gene *TBP*. * P<0.05, *** P<0.001, two-tailed unpaired t-test. **(C)** Representative H&E and IHC staining of HIF1α and GLUT1 of ccRCC and chRCC patients, scale bar is 50μm. **(D)** Quantification of HIF1α and GLUT1 staining in ccRCC and chRCC patients, categorized as intense (2), weak (1) or no detectable (0) staining. **(E)** qPCR analysis of genes involved in glycolysis and oxidative phosphorylation in chRCC tumors (n = 18) and matched kidney tissue (n = 18). Expression levels are relative to *TBP*. * P<0.05, ** P<0.01, *** P<0.001, two-tailed unpaired t-test. **(F)** Relative SF3B1 expression of the stratified TCGA-cohort. *** P<0.001, **** P<0.0001. One-way ANOVA followed by a Tukey’s multiple comparison post-test. **(G)** Survival of patients (TCGA cohort) stratified according to chromosome 2 status and SF3B1 expression as indicated. P-value was computed with a Log-rank (Mantel-Cox) test. **(H)** Percentage of chRCC patients (TCGA cohort) where spreading to lymph nodes was observed. Patients lacking information of lymph node status were omitted for analysis. Significance was computed with chi-square tests.

**Figure 6.**
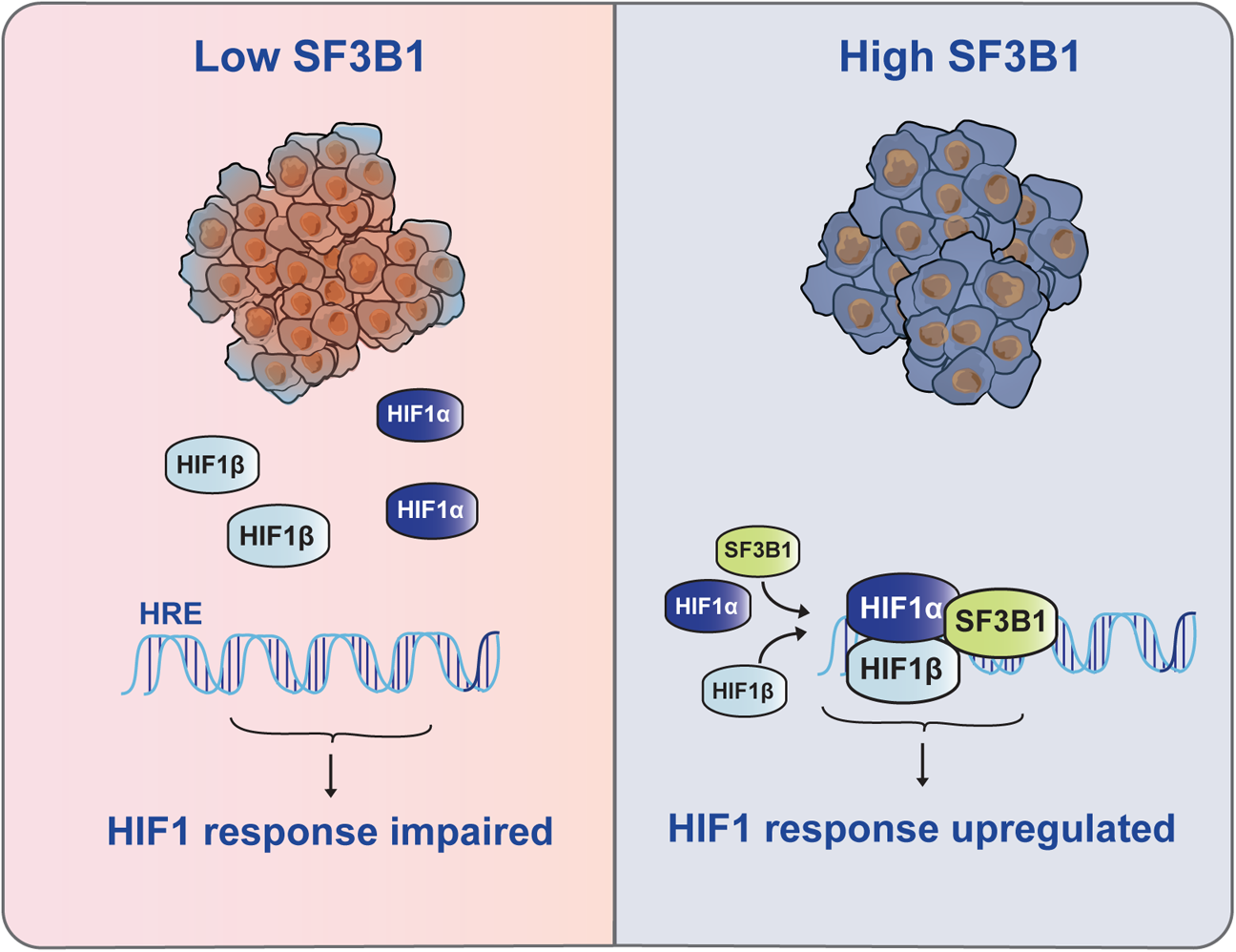
Model of HIF1 activation by SF3B1. If SF3B1 levels are low, HIF1-complex does not bind efficiently to its DNA-targets, resulting in insufficient HIF1 response and impaired metabolic adaption to hypoxia (left). When SF3B1 levels are high, a heterotrimeric HIF1 complex is formed, allowing enhanced DNA-binding and transcription of hypoxia responsive genes, resulting in metabolic adaption to hypoxia (right).

### Heterozygosity of *SF3B1* is tolerated in normoxic chromophobe renal cell carcinoma (chRCC)

Gene expression analysis in TCGA datasets, as well as knock-down experiments in melanoma- and breast cancer cell lines suggest that SF3B1-dependency for HIF target gene regulation is conserved across different solid cancers (Fig. 1A, B; Suppl. Fig. 6A-D). However, our finding that wildtype SF3B1 levels are necessary for mediating full adaptation to hypoxia suggests that frequent loss of heterozygosity in *SF3B1* should be tolerated mainly in tumors with no or mild hypoxia, which do not rely on a metabolic shift to glycolysis. To test this hypothesis, we focused on the analysis of chromophobe renal cell carcinoma (chRCC). This cancer shows an exceptional high incidence for heterozygous loss of *SF3B1*-encoding chromosome 2 (Fig. 5A), which manifests in lower *SF3B1* expression levels compared to the more frequent clear cell renal carcinoma (ccRCC) (Creighton et al., 2013; Davis et al., 2014; Ohashi et al., 2019) (Suppl. Fig. 7A). To first confirm that *SF3B1* expression also facilitates HIF signaling in chRCC, we established primary cell lines from chRCC biopsies and performed a siRNA-mediated knockdown of *SF3B1*. Reduction in *SF3B1* levels indeed led to impaired transcriptional response to hypoxia (Fig. 5B). After validating the high frequency of chromosome 2 loss in our cohort of chRCC samples (Chr2 loss in Swiss-cohort in 60% of patients - Suppl. Fig. 7B, C), we performed IHC stainings to assess HIF1α levels as a surrogate for HIF1 signaling activity. In line with our hypothesis, 90% of patient samples were negative for HIF1α-staining, with the remaining 10% exhibiting low levels (Fig. 5C, D). In contrast, in samples of clear cell renal cell carcinomas (ccRCC), a kidney cancer known for active HIF signaling(Creighton et al., 2013), moderate or high HIF1α staining was observed in 48% of samples (Fig. 5C, D). Consistent with these results, only 23% of ChRCC patients show moderate expression of the HIF target GLUT1 compared to 93% of ccRCC patients (Fig. 5C, D), and genes involved in oxidative phosphorylation, but not glycolysis, were upregulated in chRCC when compared to healthy tissue (Fig. 5E). These data support our hypothesis that loss of heterozygosity of *SF3B1* is well-tolerated in chRCC, as this cancer does not rely on a metabolic shift to glycolysis. Importantly, our data could also provide an explanation why only 4%-7% of patients of chRCC display tumor metastasis(Davis et al., 2014), as for this process adaptation to low oxygen tension is pivotal (Ohta et al., 2011). Indeed, when we stratified chRCC patients with loss of chromosome 2 into samples with low and high *SF3B1* levels (Fig. 5F; Suppl. Fig. 7D), we observed a higher incidence of lymph node spreading (7% vs 43%) and poorer prognosis in the group with upregulated *SF3B1* (Fig. 5G, H).

## DISCUSSION

Our discovery of a splicing-independent role of SF3B1 in the HIF1 complex provides an unexpected mechanistical explanation for recent studies, which correlated high SF3B1 levels with increased malignancy in solid tumors. One difficulty in our study was to detangle the role of SF3B1 in HIF-signaling and splicing. However, the 50% reduction of *SF3B1* levels resulting from loss of heterozygosity already led to a significant reduction in HIF-signaling activity without impairing splicing, and overexpression of SF3B1 led to an increase in transcriptional HIF response. In addition, while SF3B1 downregulation did not reduce HIF1 complex levels, it substantially perturbed its ability to bind to HRE regions in HIF target genes.

HIF signaling plays an essential role in most solid cancers to adapt to low oxygen environments and to induce a metabolic shift to anaerobic glycolysis. Its inhibition is therefore considered to be a highly promising treatment approach (Muz et al., 2015; Semenza, 2003; Semenza, 2019). Most HIF inhibitors tested in clinical trials, however, showed low specificity and interfered with other biological processes, leading to dose-limiting side-effects (Semenza, 2019). Since binding of SF3B1 to HIF1α is direct, inhibition of this interaction could sensitize tumor cells to low oxygen environments in a highly specific manner. Together with drugs blocking heterodimerization of HIF1α/HIF1β, such compounds might therefore enable long-term treatment without the occurrence of cross-resistance.

## ACKNOWLEDGMENTS

Prof. Wilhelm Krek (W.K., Department of Biology, Institute of Molecular Health Sciences, ETH Zurich, Switzerland), passed away in August 2018. He initiated this project and supervised the experiments until his demise. This manuscript is dedicated to his memory.

We are grateful to D. Hanahan (EPFL, Lausanne, Switzerland) and T. Jacks (Koch Institute, MIT, Cambridge/MA, USA) for providing mouse cancer cell lines and strains. We thank B. Luikart and W. Kaelin for their plasmid constructs obtained via Addgene. This work was financed by grants from the Swiss National Science Foundation, the Swiss Cancer League, the Walter Fischli Foundation, the University Research Priority Program (URPP) in Translational Cancer Research (University of Zurich) and the Fritz Thyssen Foundation. Further support was provided by the Tissue Biobank and the in-situ laboratory of the University Hospital Zurich, as well as the Molecular Genetics Laboratory at the University Children’s Hospital Zurich.

## AUTHOR CONTRIBUTIONS

P.S., C.C., P.M., W.K. and G.S. conceived the study. P.S. and C.C. designed and executed most of the experiments. G.S., W.K. and M.S. supervised the study. P.S. and G.S. wrote the manuscript with input from all authors. L.M.K. performed the experiments related to mutant SF3B1. C.H., C.P and D.L established human cancer organoid cultures. P.G. and C.M. performed isolation and cell biological characterization of murine organoids.

A.V. performed ultrasound imaging and was involved in the in-vivo study. T.R. was involved in the establishment of CRISPR-mediated SF3B1 knockout cell lines. I.G. performed in vitro-experiments. P.S., K.V.L. and Y.C. performed the *in-silico* analysis of HIF and SF3B1 expression in human cancers. N.R.D. analyzed RNA-splicing in organoids. P.S. and M.S. performed ChIP-seq analysis. G.P. performed mouse tumor tissue IHC and quantified tumor patterns. H.M., S.A. and H.A.B. performed chRCC tissue staining and evaluation and established chRCC cell lines. H.M. was responsible for diagnosis and analysis of renal tumors.

## METHODS

### TCGA data analysis

TCGA RNA-seq expression and clinical data were downloaded either from the TCGA website (https://tcga-data.nci.nih.gov), from The Human Protein Atlas (Uhlen et al., 2017) (https://proteinatlas.org) or from The cBio Cancer Genomics Portal (Cerami et al., 2017; Gao et al., 2014) (https://cbioportal.org). For survival analysis, patients were stratified by the median of *SF3B1* gene expression, except for the chRCC analysis. There, the upper quartile of *SF3B1* expression in patients with chromosome 2 loss was used for stratification. In the survival analysis of chRCC patients, a missing variable (months) of one deceased patient in the *Chr2 loss, SF3B1 high* cohort was imputed by using the mean value of all deceased patients in the unstratified dataset. The hypoxia signature was computed as the average expression of 34 hypoxia-inducible genes represented by *CA9, NDUFA4L2, EGLN3, NDRG1, PFKFB4, ADM, NRN1, CA12, ENO2, ZNF395, RAB20, DDIT4, SLC7A5, BNIP3, HMOX1, POU5F1, ALDOC, PDK1, LDHA, SLC2A1, BNIP3L, STC2, HK2, CP, LOX, MUC1, VEGFA, SLC16A3, MXI1, GYS1, COL5A1, PPP1R3C, TXNIP, LRP1*. K700E mutations are based on a uniform recalled variant dataset. RNA-seq data has been uniformly re-aligned across all 8255 cancer patient samples. Only non-alternate axons are used for gene expression quantification that has been quantile normalised (Kahles et al., 2018). Data has been made available here: https://gdc.cancer.gov/about-data/publications/PanCanAtlas-Splicing-2018.

### Cell culture

SK-MEL-28, PANC-1, MDA-MB-231, Mia-PaCa-2 and HEK293T cells were maintained in Dulbecco’s modified Eagle’s medium (DMEM, Invitrogen) supplemented with 10% FBS, and 4 mM L-glutamine. AsPC-1 and BxPC-3 cells were maintained in Roswell Park Memorial Institute (RPMI, Invitrogen) supplemented with 10% FBS and 4 mM L-glutamine. Cell lines were cultured at 37 °C in 5% CO_2_. Sf9 insect cells were grown in suspension at 27°C in Sf-900 II SFM medium (Invitrogen) supplemented with penicillin/streptomycin (Sigma-Aldrich). Cell density was monitored regularly and was maintained between 0.5 and 2 x10^6^ cells/ml. All cell lines have been authenticated by Microsynth and regular mycoplasma tests were performed using the ScienCell Mycoplasma PCR detection kit (Cat. No. 8208).

### Quantitative real-time PCR analysis

RNA was isolated using the nucleospin RNA kit (Macherey-Nagel) and reverse-transcribed to cDNA (kit: Applied Biosystems, ref. 43698813). SYBR-green based quantitative real-time PCR (qRT–PCR) was performed on a LightCycler 480 (Roche). *C*_t_ values were normalized to the housekeeping gene *TBP* for human samples and to *Actb* or *Ccny* for the mouse samples. Primer sequences are listed in Supplementary Table 3.

### Luciferase assays

Cells were co-transfected with a *Vegf*-HRE-luciferase reporter plasmid, or a *Sf3b1*-HRE-luciferase reporter plasmid together with different plasmids using lipofectamine 2000 (Invitrogen). A renilla luciferase plasmid was co-transfected for internal control. Cells were collected 24 hrs after transfection, and luciferase activity analyzed on a TECAN M1000 PRO using the dual-luciferase reporter assay system (Promega E1980). The assays were carried out according to the manufacturer’s instructions.

### Transient transfections

siRNAs were transfected using Lipofectamine-RNAiMax reagent (Invitrogen). We used Qiagen human si*SF3B1*#1 (SI04159456), human si*SF3B1*#2 (SI04161766) and human si*HIF1Α 5’-*CUG AUG ACC AGC AAC UUG A-3’ and 5’-UCA AGU UGC UGG UCA UCA G-3’. For transient overexpression studies, DNA plasmids were transfected with Lipofectamine 2000 (Invitrogen) according to the manufacturer’s protocol.

### Lentivirus production and transduction

Lentiviral particles were produced in HEK-293T cells. HEK-293T cells were purchased from American Type Culture Collection (ATCC). Cells were transduced with lentivirus by adding virus-containing supernatants to the cell culture medium.

### Chromatin immunoprecipitations (ChIP)

ChIP assays were performed using the ChIP-IT Kit (Active Motif) according to the manufacturer’s instructions. ChIP was performed with HIF1*α* antibody from Abcam (ab2185), SF3B1 antibody from MBL International (D221-3), an HIF1β antibody from Novus Biological (NB 100-124), or an anti-IgG antibody from Abcam (ab171870). Primer sequences used are listed in Supplementary Table 3.

### ChIP-sequencing: Sample preparation

HIF1α-ChIP-assays for ChIP-seq was performed using the SimpleChIP® Enzymatic Chromatin IP Kit (#9003) from Cell Signalling Technologies according to the manufacturer’s instructions. In brief, cells from 4 15-cm dishes were cross-linked using formaldehyde for 5 minutes under hypoxia, the cross-linking was subsequently quenched with incubation of with 125 mM glycine (final concentration) for 5 minutes at normoxia and the cells were collected. Cells were lysed and nuclei were isolated by centrifugation. DNA was fragmented by micrococcal nuclease digestion for 20 minutes at 37°C, and nuclei were disrupted by brief sonication 10 minutes on a high setting. Cleared extracts were subjected for immunoprecipitation using HIF1α (D1S7W XP® Rabbit mAb #36169) and IgG antibodies as control. After reversal of the cross-link, DNA was isolated and its concentration measured with Qubit dsDNA HS assay kit (Q32851) according to the manufacturers instruction. For controlling potential amplification-biases during library-preparation, 0.2 ng of drosophila chromatin, treated as described above for the experimental samples, was spiked-in for every sample before library-preparation.

### ChIP-sequencing: Library-preparation and sequencing

The NEBNext® Ultra™ II DNA Library Prep Kit from NEB (New England Biolabs, Ipswich, MA) in the succeeding steps. ChIP samples (1 ng) were end-repaired and adenylated. Adapters containing the index for multiplexing were ligated to the fragmented DNA samples. Fragments containing adapters on both ends were enriched by PCR. The quality and quantity of the enriched libraries were measured using Qubit® (1.0) Fluorometer and the Tapestation (Agilent, Waldbronn, Germany). The product is a smear with an average fragment size of approximately 300 bp. The libraries were normalized to 10nM in Tris-Cl 10 mM, pH8.5 with 0.1% Tween 20. The libraries were sequenced single read 100 bp using the Illumina Novaseq 6000 (Illumina, Inc, California, USA). Reads were quality-checked with fastqc which computes various quality metrics for the raw reads.

### ChIP-sequencing: Computational analysis

SnakePipes “ChIP-seq” pipeline (Bhardwaj et al., 2019) was used for the analysis of the sequencing data according to the developer’s guidelines. In brief, reads were trimmed, deduplicated and aligned simultaneously to the masked genomes of homo sapiens (GRCh38.p13) and drosophila melanogaster (BDGP6.32) published by Ensembl. Three replicates for each condition were used. On average, 86 million (SD: 18 million) reads per sample were uniquely mapped to the hybrid genome. Peaks were identified by MACS2(Zhang et al., 2008), using merged reads of two chromatin inputs per condition for background normalization. CSAW (Lun and Smyth, 2015) was applied for differential binding analysis, whereby size factors were calculated by computing the relative number of mapped reads to the drosophila genome for each sample. Genes in proximity of peaks were identified by searching for nearest transcripts. For subsequent analyses, only peaks that were identified in all of the 3 siCtrl-ChIP replicates were considered (high-confidence peaks). Motif recognition was performed with MEME (Bailey and Elkan, 1994), using sequences of −49 and +50 bp of the summit of the respective peak. A maximum of 1000 input sequences was considered for the motif-recognition. The following parameters were used: *-dna -mod zoops-objfun cd -revcomp -minw 10 -maxw 15*. The heatmap of high-confidence peaks was computed with deeptools/compute matrix and /plotheatmap. Exemplary peaks were visualized using pyGenomeTracks on two corresponding replicates with comparable sizefactors (0.93 / 0.91) to facilitate visual comparison. Pathway analysis on the gene-set described in Fig. 3F & G was performed with Enrichr on the BioPlanet database.

### Subcellular protein fractionation

Subcellular protein fractionation and isolation of chromatin fractions was performed using the Subcellular Protein Fractionation Kit for Cultured Cells kit from Thermo Fisher, following the instructions of the manufacturer. Soluble nuclear and chromatin-bound nuclear extracts were loaded on an SDS–PAGE gel and subjected to western blotting.

### Immunoblotting and antibodies

Protein lysates were resolved on polyacrylamide minigels and transferred onto nitrocellulose membrane by semi-dry transfer. The following antibodies were used for immunoblotting: rabbit HIF1*α* (catalogue no. NB-100-479; Novus Biologicals), mouse SF3B1 (catalogue no. D221-3; MBL International), mouse HIF1α (catalogue no. NB 100-124; Novus Biologicals), mouse SF3B2 (catalogue no. ab56800; Abcam), rabbit SF3B4 (catalogue no. ab157117; Abcam), rabbit SF3B14 (catalogue no. NBP1-87431; Novus Biologicals), rabbit α-tubulin (catalogue no. ab18251; Abcam), rabbit CA9 (catalogue no. NB-100-417; Novus), rabbit PKM2 (catalogue no. 3198; Cell Signaling), rabbit PDK1 (catalogue no. 3061S; Cell Signaling), mouse HA-tag (catalogue no. 26183; Thermo Scientific), rabbit p53 (catalogue no. FL-393; Santa Cruz), mouse KrasG12D (catalogue no. 26036; Neweastbio), mouse total-Ras (catalogue no. 3965; Cell Signaling), rabbit p-Mek (catalogue no. 9154; Cell Signaling), rabbit total-Mek (catalogue no. 9122S; Cell Signaling), rabbit p21 (catalogue no. ab109199; Abcam), rabbit p16 (catalogue no. ab51243; Abcam). Immunodetection and visualization of signals by chemiluminescence was carried out by using the ImageQuant LAS 4000 mini (GE Healthcare). For immunoprecipitation, transfected cells were lysed in cell lysis buffer TNN (50 mM Tris-HCl pH 7.5, 250 mM NaCl, 5 mM EDTA, 0.5% NP-40, 50 mM NaF, 1 mMDTT, 1 mM PMSF, and protease inhibitor cocktail) for 30 min. Magnetic dynabeads (Invitrogen) were incubated first with the primary antibody for 10 min at room temperature, washed and then incubated with the cell lysate for 3 hours at 4°C. The assays were carried out according to the manufacturer’s instructions. Finally, the beads were boiled in 2x sample buffer for 5 min and eluents were analyzed by Western blotting. For co-immunoprecipitation with recombinant proteins, magnetic dynabeads (Invitrogen) were incubated first with the primary antibody and then incubated with the recombinant proteins for 2 hours at 4°C.

### Human PDAC organoid culture

Human PDAC organoids were established from patients’ surgical specimens and cultured as described in full detail by (Hirt *et al*, manuscript under revision). Human pancreatic tissue samples were provided by the Department of Pathology and Molecular Pathology, University Hospital Zurich, based on informed consent and study approval from the ethical committee (BASEC-Nr. 2017-01319). For all samples, the diagnosis of PDAC was confirmed on corresponding tissue slides reviewed by board-certified pathologists. To establish organoid lines, tissue was chopped and digested in full medium containing collagenase type II (5 mg/ml). The digestion was stopped with advanced DMEM/F12 medium supplemented with HEPES (10mM), Glutamax (1%) and penicillin/streptomycin (1%). Cells were seeded as 20μl drops of Matrigel (Corning, growth factor reduced) into a 48-well suspension culture plate. Human PDAC organoids were cultured in Advanced DMEM supplemented with 10 × 10^−3^ M HEPES, 1x Glutamax, 1% Penicillin/Streptomycin, 1x B27, 1.25 × 10^−3^ M N-acetylcysteine, 50% Wnt3a conditioned medium (CM), 10% R-spondin-1 CM, 10% noggin CM, 10 × 10^−3^ M nicotinamide, 1 × 10−6 M prostaglandin E2, 50 ng mL^−1^, EGF, 10 × 10^−9^ M gastrin, 100 ng mL^−1^ FGF10 and 0.5 × 10^−6^ M A83-01.

### RCC ethics statement and patient selection

All tissue samples were made available by the Tissue Biobank of the Department of Pathology and Molecular Pathology, University Hospital of Zurich, Switzerland and collected as described previously (Bolck et al., 2019). The local ethics commission approved this study (BASEC_2019-01959) and patients gave written consent. The retrospective use of normal and tumor tissues of RCC patients is in accordance with the Swiss Human Research Act, which allows the use of biomaterial and patient data for research purposes without informed consent under certain conditions that include the present cases (article 34). All tumors were reviewed by a board-certified pathologist and histologically classified according to the World Health Organization guidelines.

### ChRCC tissue processing and generation of cell models

Renal tumors were surgically resected and underwent routine tissue processing and rapid sectioning for diagnostic purposes and generation of cell models (including formalin fixation and paraffin embedding, snap freezing of fresh tumor and normal tissue). Fresh tissue samples macroscopically identified as cancer by a pathologist with specialization in uropathology (H. Moch) were placed into sterile 50-ml conical tubes containing transport media (RPMI (Gibco, Waltham, MA) with 10 % fetal calf serum (FCS, Gibco) and Antibiotic-Antimycotic® (Gibco)) and stored at 4 °C until further processing. Briefly, samples were rinsed with PBS and subsequently cells were isolated by finely cutting the tissue into small pieces followed by collagenase A digestion for 1-3 hours at 37 °C. The slurry was passed through a 100 µm cell strainer to remove large fragments and the Collagenase A digestion was quenched with DMEM containing 10-20 % FCS. Cells were washed once with PBS and if necessary, erythrocytes were lysed. Finally, cell viability was evaluated by trypan blue dye exclusion and an appropriate number of cells was resuspended in Renal Epithelial Cell Growth Medium 2 (Promocell). This suspension was seeded into collagen I-coated cell culture dishes (Corning, Corning, NY) containing 30–50 % confluent Mitomycin C-treated mouse embryonic fibroblasts CF6 (ThermoFisher). Subsequently, the co-culture was maintained in Renal Epithelial Cell Growth Medium 2 in a humidified incubator at 37 °C with 5% CO_2_. The medium was replaced earliest 5 days after initial plating and subsequently every three to four days. Cells were expanded by passaging without the addition of new CF6 feeder cells in subsequent passages.

### RCC immunohistochemistry

For immunohistochemical (IHC) analyses, a tissue microarray (TMA) and whole tissue sections of primary RCCs from the Department of Pathology and Molecular Pathology, University Hospital of Zurich, Switzerland were used (Dahinden et al., 2010). Clinicopathological parameters of the RCC specimens are described in Supplementary Table 4. Formalin-fixed paraffin-embedded (FFPE) cell pellets from cell cultures were prepared as previously described (Struckmann et al., 2008). For histological evaluation, FFPE specimens were sectioned (2 µm) and stained with hematoxylin and eosin (HE). For IHC, 2 μm sections were transferred to glass slides followed by antigen retrieval. Antibodies used for IHC are summarized in the table below. IHC was performed using the Ventana Benchmark automated system (Roche, Ventana Medical Systems, Oro Valley, AZ) and Ventana reagents. The Optiview DAB IHC detection and the Discovery Chromomap DAB Kits (Roche, Ventana Medical Systems) were used to stain with the antibodies against Glut1 and SF3B1, respectively. The staining procedure for HIF1α was carried out with the automated Leica BOND system using the Bond Polymer Refine Detection Kit (Leica Biosystems, Wetzlar, Germany).

| Antibody | Name/Clone | Dilution IHC | Supplier |
| --- | --- | --- | --- |
| SF3B1 | EPR11986 | 1:100 | Abcam |
| HIF1α | HIF-1 alpha antibody [mgc3] | 1:400 | Abcam |
| GLUT1 | Glucose Transporter type 1 | 1:1000 | Millipore |

### RNA-extraction of chRCC patient samples

RNA extraction from FFPE specimens of chRCC was performed using the Maxwell® 16 Tissue DNA Purification Kit (Promega). RNA quality was assessed with the RNA Qubit™ RNA HS Assay Kit (ThermoFisher). Subsequently, cDNA was prepared using the Superscript IV Vilo Mastermix (Thermo Fisher).

### SF3B1 quantification in chRCC patient samples

The quantitative measurements of SF3B1 were performed using the Taqman Fast Advanced master mix (Thermo Fisher) with 5 ng/μL ng of cDNA in each technical duplicate and the cycling parameters according to the protocol on a ViiA7 (Thermo Fisher). Primer and probe set assay IDs for the TaqMan assays were Hs00961640_g1 for SF3B1 and Hs03929097 for GAPDH, (ThermoFisher). Normal tissue was used to normalize the quantitative analysis of all samples. For the other genes assessed, regular qPCR was performed as described above.

### Analysis of Copy Number Variations (CNVs)

DNA was processed and hybridized to Affymetrix CytoScan HD array according to manufacturer’s protocol. Sample level quality control was initially assessed by Affymetric ChAS software, version 2.0 and probeset.txt files were exported. These served as input for the *rCGH* R package (v. 3.8) obtained from Bioconductor. The comprehensive comparative genomic hybridization (CGH) analysis workflow provided by *rCGH* was used for log2 relative ratio (LRR) calculation (the sample DNA signals against a normal two-copy DNA reference), profile centralization and profile segmentation. Significant focal copy number alterations were identified from segmented data using GISTIC 2.05. Subsequently, absolute copy numbers were inferred.

### Plasmid design

Deletion mutants of HIF1*α* were generated by PCR using a pcDNA3-HA-HIF1*α* plasmid (addgene 18949) as template. We designed specific primers that border the domain to be deleted on both sides in order to remove the different targeted domains of HIF1*α*.

### Baculovirus expression system

Baculovirus expression vectors expressing Strep-Tag II epitope tagged-SF3B1, HIF1α, or HIF1β were constructed by inserting the cDNA of corresponding genes into a pFBDM plasmid. Strep-Tag II tagged-SF3B1, HIF1α, or HIF1β were expressed in Sf9 cells that had been infected with recombinant baculovirus as described in the Bac-to-Bac® Baculovirus Expression System from Invitrogen.

### Immunoprecipitation and Co-immunoprecipitation (CoIP) assays

For immunoprecipitation, transfected cells were lysed in cell lysis buffer TNN (50 mM Tris-HCl pH 7.5, 250 mM NaCl, 5 mM EDTA, 0.5% NP-40, 50 mM NaF, 1 mM DTT, 1 mM PMSF, protease inhibitor cocktail and Pierce™ Universal Nuclease for Cell Lysis) for 30 min. Magnetic dynabeads (Invitrogen) were incubated first with the primary antibody for 10 min at room temperature, washed and then incubated with the cell lysate for 3 hours at 4°C. The assays were carried out according to the manufacturer’s instructions. Finally, the beads were boiled in 2x sample buffer for 5 min and eluents were analyzed by Western blotting. For co-immunoprecipitations with transiently overexpressed flag-Sf3b1 and flag-Sf3b1^K700E^, transfected HEK293T cells were lysed in TNN buffer as described above. Lysates were cleared by centrifugation at 16 000 x g for 30 minutes. Anti-FLAG® M2 Magnetic Beads (Merck) were equilibrated in lysis buffer and incubated with the cell lysate for 1 hours at 4°C. The beads were washed at least 4 times with TNN buffer and boiled in 2x sample buffer for 5 min. The eluents were analyzed by western blotting. For co-immunoprecipitation with recombinant proteins, magnetic dynabeads (Invitrogen) were incubated first with the primary antibody and then incubated with the recombinant proteins for 2 hours at 4°C.

### Mouse strains

*LSL-Kras^G12D/+^; LSL-Trp53^R172H^* (KP), and *LSL-Kras^G12D/+^; LSL-Trp53^R172H/+^; Ptf1a-Cre* (KPC) were obtained from Tyler Jack’s (Howard Hughes Medical Institute). Sf3b1 flox/flox mice were generated by targeting exon five and six of the Sf3b1 gene. These exons were floxed with LoxP sites and the targeted region was about 3.0 kb. A LoxP/FRT-flanked neomycin resistance cassette was placed 277 bp upstream of exon five and six for selection. Chimeric mice were generated via electroporation in embryonic stem cells followed by microinjection in blastocysts and implantation into foster mice. For all *in vivo* experiments, except the survival experiment, mice between the age of 7 and 13 weeks were used. Female and male mice were used for all experiments. All mice were maintained in a SPF animal facility at the ETH Phenomics Center EPIC at ETH Zürich. Maintenance and animal experiments were conducted in accordance with the Swiss Federal Veterinary Office (BVET) guidelines. Animal surgeries and ultrasound (Vevo 2100 system) analysis was performed blinded by a veterinarian. Animal numbers for experiments were chosen based on expected mortality rates and phenotypical changes of the pancreas in mice.

### Histological and immunohistochemical analysis (murine specimen)

Following euthanasia, pancreata were removed, weighed and fixed overnight in 10% neutral buffered formalin (Formafix Switzerland AG, Hittnau, Switzerland). The tissue was embedded in paraffin and cut serially into 3-5-μm sections, which were stained with hematoxylin and eosin (HE) and Masson trichrome for the assessment of fibrosis. Histological analysis, including murine pancreatic intraepithelial neoplasia (mPanIN) grading and assessment of the proportion of affected versus unremarkable pancreatic parenchyma, was carried out in the HE-stained sections in a blinded fashion by an ECVP-board certified veterinary pathologist (GP), according to previously reported criteria(Hruban et al., 2006). Immunohistochemistry (IHC) was conducted on consecutive paraffin-embedded sections using the primary antibodies against HIF1α (rabbit polyclonal, Novus Biological, NB100-479, 1:500), SF3B1 (rabbit monoclonal, clone number EPR11986, Abcam, ab172634, 1:100), GLUT-1 (rabbit polyclonal, Merck Millipore, #400060, 1:100), Ki-67 (rabbit monoclonal, clone name 30-9, Ventana Roche, 790-4286, 1:400), cleaved caspase 3 (rabbit monoclonal, clone number D3E9, Cell Signaling, Cell Signaling, #9579, 1:400) and CD31 (rabbit polyclonal, Santa Cruz, sc-1506R, 1:1000). All immunostains were performed in Ventana Discovery (Ki-67 and cleaved caspase 3) or Dako autostainers using 3,3’ diaminobenzidine (DAB) chromogen (Dako-Agilent Technologies, Denmark). All slides were scanned using digital slide scanner NanoZoomer-XR C12000 (Hamamatsu, Japan) and images were taken using NDP.view2 viewing software (Hamamatsu). mPanIN were counted manually in 10 high power fields (HPF) in each HE-stained section. Automated quantitative analysis was carried out on the digital slides using the Visiopharm Integrator System (VIS, version 4.5.1.324, Visiopharm, Hørsholm, Denmark). Briefly, a linear threshold classification allowed recognition of the blue (Masson trichrome) or DAB brown (IHC) structures in 20 regions of interest (ROI) sized 0.237 mm^2^ (the area of an HPF with one ocular of 22 mm field of view), randomly selected across the pancreatic parenchyma. Results were expressed as percentage of positive area/ROI vs total parenchyma/ROI (Masson trichrome, HIF-1*α*, GLUT1, CD31) or average number of positive nuclei or cells/ROI (SF3B1, Ki-67, cleaved caspase 3).

### Heterozygous knockout of SF3B1 by CRISPR-Cas9

We used the sgRNA-Designer platform by the Broad Institute to design sgRNAs for targeted gene knock-out (https://portals.broadinstitute.org/gpp/public/analysis-tools/sgRNA-design). Two distinct sgRNAs were used: sgSF3B1_A (5’-TAATCTTCATCAATCAATAG-3’) and sgSF3B1_B (5’-AAGATCGCCAAGACTCACGA-3’). According to the protocol from Shalem et al., we adapted the sgRNA design for subsequent cloning into the LentiCRISPRv2 backbone (Shalem et al., 2014). Cells were transduced with lentivirus and selected with puromycin. Heterozygous knock-out of the target gene was validated with deep sequencing for PANC-1 and AsPC-1 cells at two distinct time-points, to assess the enrichment of in-frame indels. Primers were designed flanking the predicted sgRNA cut-site and containing primer binding sites for Illumina TruSeq Deep sequencing primers. After purification, the amplicons were sequenced on an Illumina MiSeq System, with a minimum of 5’000 reads. Analysis of editing events was performed with CRISPResso V1.0.1.

### Murine pancreatic ductal organoid culture

Pancreatic ducts were isolated from the whole organ of 13 weeks old KP and KPS mice as previously described (Huch et al., 2013). Each organoid line was isolated from an individual mouse. Isolated ducts as well as the ensuing organoids were embedded in growth-factor reduced (GFR)-Matrigel (Corning), and cultured in organoid medium (OM), which is composed of AdDMEM/F12 (Gibco) supplemented with GlutaMAX (Gibco), HEPES (Gibco), Penicillin-Streptomycin (Invitrogen), B27 (Gibco), N-2 (Gibco), 1.25 mM N-Acetyl-L-cysteine (Sigma), 10 nM Gastrin I (Sigma) and the growth factors: 100 ng/ml FGF10 (Peprotech), 50 ng/ml EGF (Peprotech), 100 ng/ml Noggin, 100 ng/ml RSPO-1 (Peprotech), and 10 mM Nicotinamide (Sigma). For the first week after duct isolation the culture medium was supplemented with 100 µg/ml Primocin (InvivoGen).

### Lentiviral production and transduction of organoids

The plasmids FUG-T2A-GFP-Cre and control GFP were purchased from Addgene #66691 and lentiviral particles were produced in HEK-293T cells using X-tremeGene 9 DNA Transfection Reagent (Roche). Upon ultracentrifugation concentrated virus particles were stored at −80°C and used to infect pancreatic ductal organoid cultures as previously described (Huch et al., 2013). Briefly, upon dissociation of the GFR-Matrigel, organoid cultures were broken down into single cells using TrypLE Express (Gibco) and fire-polished glass Pasteur pipettes. Single cells were transferred to maximum recovery microtubes (Axygen) and centrifuged at 800 g for 5 min at 4°C. The pellets were resuspended in 250 µl OM and inoculated with 10 µl of concentrated virus particles that was then incubated at 37°C for 4-6 h. After a washing step, the cells were seeded in GFR-Matrigel and grown in OM. Two to three weeks later the pool of GFP-expressing organoids was dissociated again into single cells and subjected to FACS analysis (MoFlo® Astrios™ EQ) to sort for GFP-positive cells. The cells were put back into culture and used for subsequent experiments.

### Organoid proliferation assay

Organoids were seeded as single cells at a density of 500 cells/μl GFR-Matrigel (Corning). 48 hours after seeding, cells were exposed to 1% O_2_ in a hypoxia chamber (SCI-tive Hypoxia Workstation, Baker Ruskinn) or kept in a cell culture incubator at 21% O_2_. At depicted time-points, the organoids were recovered from the matrigel and dissolved in 200 μl TE-buffer. After 3 freeze-thaw cycles, the amount of DNA was quantified with Quant-iT™ PicoGreen™ dsDNA Assay Kit (ThermoFisher Scientific) according to the manufacturer’s instructions. Absolute values were normalized to the DNA-content 48 hours after seeding.

### [^14^C] 2-Deoxy-D-glucose uptake

For organoid experiments, 3 wells of 50 µl matrigel-embedded organoids per condition were seeded in a 24-well plate. On day 3, cells were kept in normoxia or exposed to hypoxia (at 1% O_2_) for 12 hours. The medium was replaced with DMEM containing uniformly-labelled [^14^C] 2-Deoxy-D-glucose (Hartmann Analytic) at a concentration of 150 μci/mmol and 5 mM glucose. Cells were washed twice with PBS just before harvesting. Radioactivity counts were normalized to cell number.

### Illumina RNA sequencing experiment

Multiple organoid lines (5x KPC, 4x KPCS) were seeded as described, where each line was derived from an individual mouse. After 48 hours of exposure to 1% O_2_ or 21% O_2_, organoids were harvested and RNA was extracted using the NucleoSpin^®^ RNA kit (Macherey-Nagel) according to the manufacturer’s instructions. For each organoid line, 2-3 wells of organoids from a 24 well plate were harvested in lysis buffer RA1 (RNA extraction kit, MN) supplemented with TCEP (Tris(2-carboxyethyl)phosphine hydrochloride). For library preparation, the quantity and quality of the isolated RNA was determined with a Qubit® (1.0) Fluorometer (Life Technologies, California, USA) and a Bioanalyzer 2100 (Agilent, Waldbronn, Germany). The TruSeq Stranded mRNA Sample Prep Kit (Illumina, Inc, California, USA) was used in the succeeding steps. Briefly, total RNA samples (1 μg) were ribosome depleted and then reverse-transcribed into double-stranded cDNA with Actinomycin added during first-strand synthesis. The cDNA samples were fragmented, end-repaired and polyadenylated before ligation of TruSeq adapters. The adapters contain the index for multiplexing. Fragments containing TruSeq adapters on both ends were selectively enriched with PCR. The quality and quantity of the enriched libraries were validated using Qubit® (1.0) Fluorometer and the Bioanalyzer 2100 (Agilent, Waldbronn, Germany). The product is a smear with an average fragment size of approximately 360 bp. The libraries were normalized to 10nM in Tris-Cl 10 mM, pH8.5 with 0.1% Tween 20. Cluster generation and sequencing involved application of the TruSeq SR Cluster Kit v4-cBot-HS or TruSeq PE Cluster Kit v4-cBot-HS (Illumina, Inc, California, USA) using 8 pM of pooled normalized libraries on the cBOT. Sequencing were performed on the Illumina HiSeq 4000 paired end at 2x 100 bp (2x at least 40 million reads) using the TruSeq SBS Kit v4-HS (Illumina, Inc, California, USA).

### RNAseq data analysis

The fastq data for the read pairs from the Illumina sequencer has been processed in a bioinformatic workflow. First, the remains of the adapters and sub-standard quality reads have been trimmed with trimmomatic (v0.35). For the pairs in which both reads have passed the trimming, they have been mapped to the respective genome using STAR (v2.7.0a) and indexed BAM files obtained with samtools (v1.9). The reads have been counted using featureCounts from subread package (v1.5.0) with the options for pairs of reads to be counted as RNA fragments. The counting used Ensembl v95 mouse genome annotation GTF file. The count vectors for all the samples have been combined into a count table. The table has been processed in the secondary (statistical) analysis with R scripts using edgeR (v3.24.3), in particular binomial generalized log-linear model with contrast tests. It resulted in lists of genes ranked for differential expression by p-value and used Benjamini-Hochberg adjusted p-value as the estimate of the false discovery rate. Read counts are listed in Supplementary Table 5.

### Splicing analysis

All splice events were identified using SplAdder on the aligned RNA-Seq BAM files (Kahles et al., 2016). Gencode release M20 (Mus_musculus.GRCm38.95) was used for identification of slice events. Splice events considered were exon skips, alternative 3’/5’, and intron retentions. Only events with highest confidence as defined by SplAdder (level 3) were considered for downstream analyses. All parameters used for the analyses are listed here:

*-v y -M merge_graphs -t exon_skip,intron_retention,alt_3prime,alt_5prime -c 3 --ignore_mismatches y*

Only events with a minimum of 5 reads and an average expression greater than 20 reads in the flanking exons across all samples were considered in the analysis. All plots and analyses performed on SplAdder output were done in R (version 3.3.3). Intron retention events are listed in Supplementary Table 6.

### Statistical analysis

Statistical significance was determined using the statistical tests indicated in the respective figure legends. Data are presented as mean ± SD, if not indicated differently. Mouse studies were not randomized, but were blinded as the analyses were conducted on randomly assigned mouse number, and given to operators without any information until the end of the experiments. Animals of both genders were used and mice with health concerns were excluded from experiments. No inclusion/exclusion criteria were pre-established. No statistical methods were used to predetermine sample size. All experiments are performed at least 3 times if not indicated differently; n refers to the number of biological replicates.

---

**Supplementary Figure 1.**
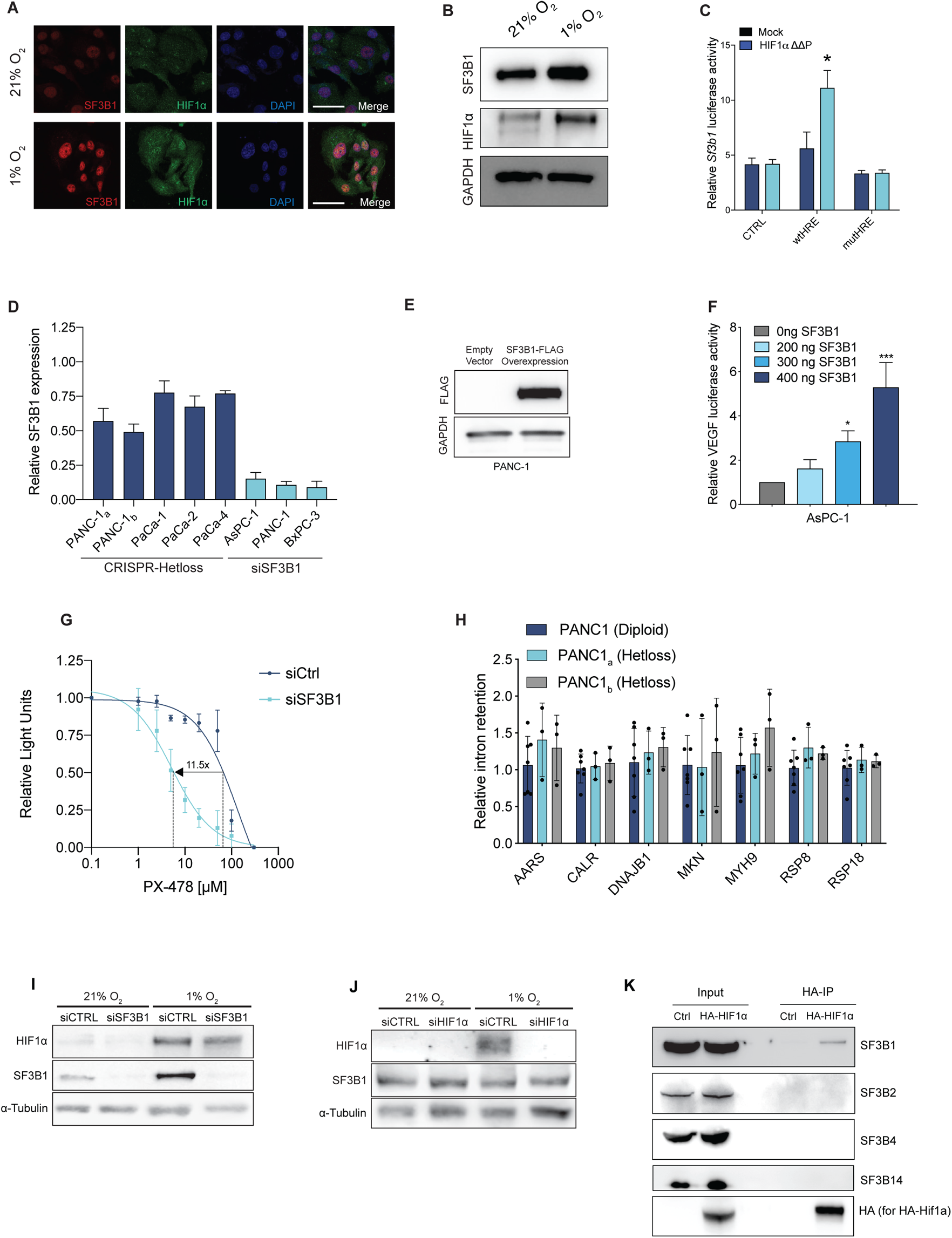
Effects of SF3B1 reduction on HIF-response. **(A)** Immunofluorescence images of PANC-1 cells grown in 21% O_2_ or 1% O_2_ and stained for the indicated proteins. **(B)** Western blot of SF3B1, HIF1α and GAPDH of PANC-1 cells exposed to 21% or 1% O_2_ for 12 hours. **(C)** Transfection of a wildtype (WT) or HRE-mutated *SF3B1* promoter-luciferase reporter (mutHRE = AAAAA) together with either an empty vector control, or HIF1αΔΔP in SK-MEL28 cells. Data are shown as mean ± S.D. of biological replicates (n=3). ** P<0.01 normalized to mock wtHRE, two-tailed unpaired t-test. **(D)** Relative SF3B1expression in the indicated patient-derived organoids and cell lines, engineered with CRISPR-Cas9 (mono-allelic loss of *SF3B1*) or treated with siRNA, and measured by qPCR. In PANC-1 cells two different guide RNAs were used to target *SF3B1*. Data are shown as mean ± S.D. of biological replicates (n = 3). The data is normalized to the respective unedited cell line (CRISPR-Hetloss) or treatment with control siRNA. Expression levels are relative to *TBP*. **(E)** Western blot analysis for FLAG in protein extracts from PANC-1 cells overexpressing FLAG-tagged SF3B1 **(F)** Transfection of a *VEGF* promoter-luciferase reporter alone or together with either HIF1αΔΔP (the constitutively active form of HIF1α), or in combination with different concentrations of SF3B1 plasmid in AsPC-1 cells. **(G)** VEGF promoter-luciferase reporter activity in PANC-1 cells treated with control siRNA or *SF3B1*-targeting siRNA (siSF3B1) cultured in hypoxia for 24 hours and exposed to the indicated doses of PX-478 (n=4). **(H)** Intron retention assessed by qPCR; expression levels are relative to the housekeeping gene *TBP*. Data are shown as mean ± S.D. of biological replicates (n ≥ 3). One-way ANOVA followed by a Dunnet’s multiple comparison post-test. **(I and J)** Protein extracts from HEK293T cells treated with siRNA targeting SF3B1 (G) or HIF1α (H), exposed to 21% or 1% O_2_ for 12 hours and processed for immunoblotting for the indicated proteins. **(K)** PANC-1 cells were transfected with HA-tagged HIF1α and cultured in hypoxia. Lysates were then processed for immunoprecipitation with an anti-HA antibody (HA-IP) followed by immunoblotting for the indicated proteins.

**Supplementary Figure 2.**
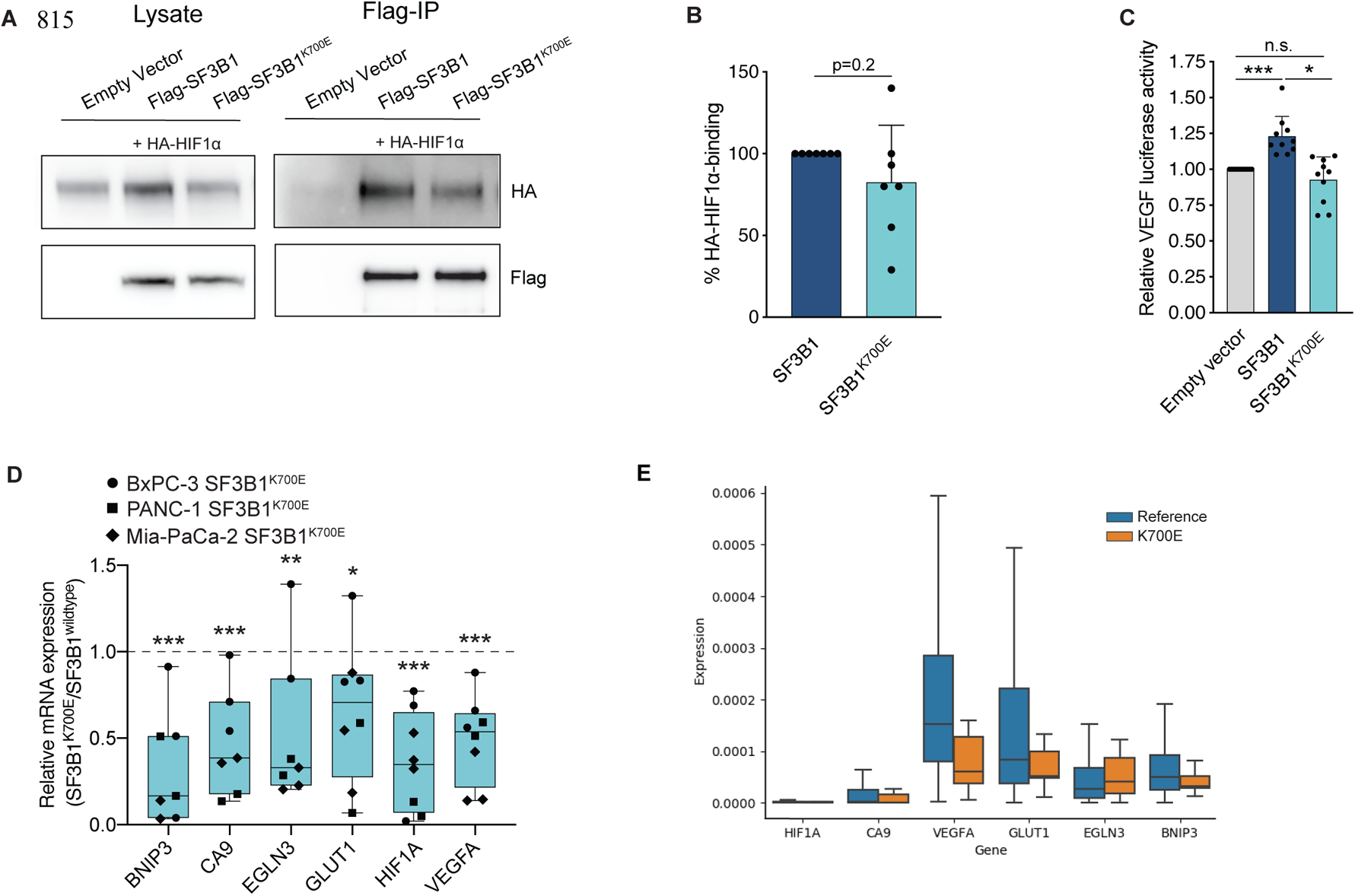
SF3B1-K700E mutation does not increase HIF1-response. **(A)** Representative Western-blot of the Co-IP of HA-tagged HIF1α and FLAG-tagged wildtype SF3B1, and its variant K700E. **(B)** Quantification of HA-tagged HIF1α in HEK293T cells, pulled down with the indicated SF3B1 variants. The data are represented as mean ± S.D. of independent experiments (n = 7). P-value was calculated by a two-tailed unpaired t-test. **(C)** Transfection of the indicated constructs in HEK293T cells stably expressing a HIF-luciferase reporter. Data are shown as mean ± S.D. of biological replicates, indicated by data points. *<P0.05, **P<0.01, ***P<0.001, two-tailed unpaired t-test. Data are normalized to transfection of an empty vector construct. **(D)** qPCR analysis of HIF-target genes in the indicated cell lines overexpressing *SF3B1^K700E^* after exposure to 1% O_2_ for 8 hours. Data are normalized to the respective cell lines overexpressing wildtype *SF3B1* after exposure to 1% O_2_ for 8 hours. Expression levels are relative to the housekeeping gene *TBP* (n = 2-3). Each data point represents an individual experiment of the depicted cell line. * P<0.05, ** P<0.01, *** P<0.001, two-tailed unpaired t-test. **(E)** Pan-cancer analysis of selected HIF-target genes in patient samples with *SF3B1^K700E^* (orange) or *SF3B1* (blue). Only solid cancers in the TCGA database were used for analysis (reference = 8243 samples; *SF3B1* = 12 samples).

**Supplementary Figure 3.**
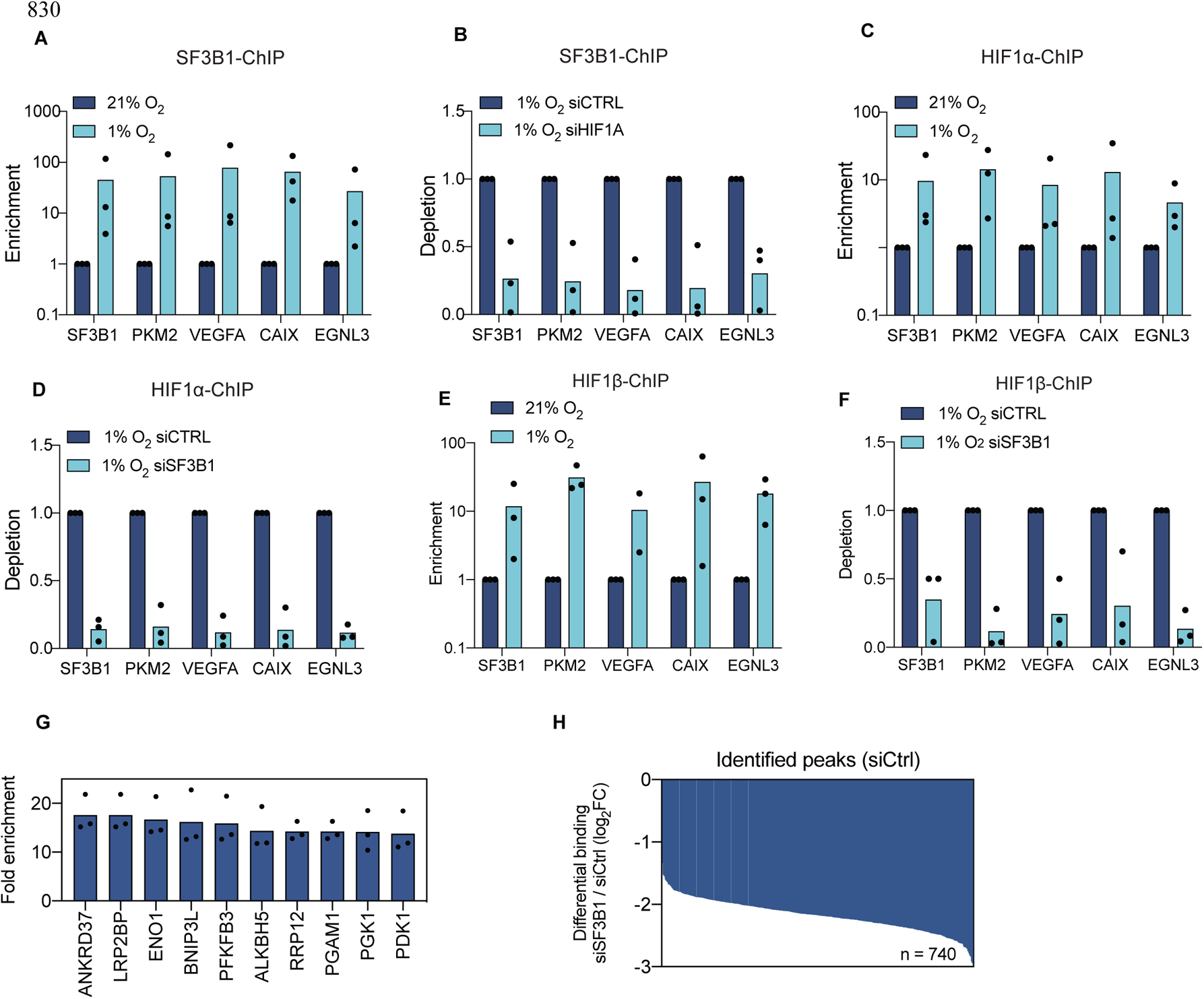
SF3B1 increases binding of HIF1α to its target genes. **(A to F)** Chromatin immunoprecipitation (ChIP) using a specific antibody against SF3B1 (A, B) HIF1α (C, D) or HIF1b (E, F) in HEK293T cells. (A, C, E): Cells are cultured under 21% or 1% O_2_ for 8 hours. The fold-increase in target gene binding under hypoxia is shown relative to normoxia. (B, D, F): Cells are transfected with siRNA targeting HIF1α (B), SF3B1 (D, F) or scrambled siRNA as control, and incubated in 1% O_2_ for 8 hours. The decrease in target gene binding in siRNA treated cells is shown relative to the siRNA control. Results show three independent experiments. **(G)** Fold enrichment over input of the 10 highest enriched peaks of the siCtrl-treated samples identified by HIF1α-ChIP-Seq. **(H)** Differential binding analysis of peaks shown in Fig. 3D in PANC-1 cells treated with siSF3B1 compared to siCtrl, identified by HIF1α-ChIP-Seq.

**Supplementary Figure 4.**
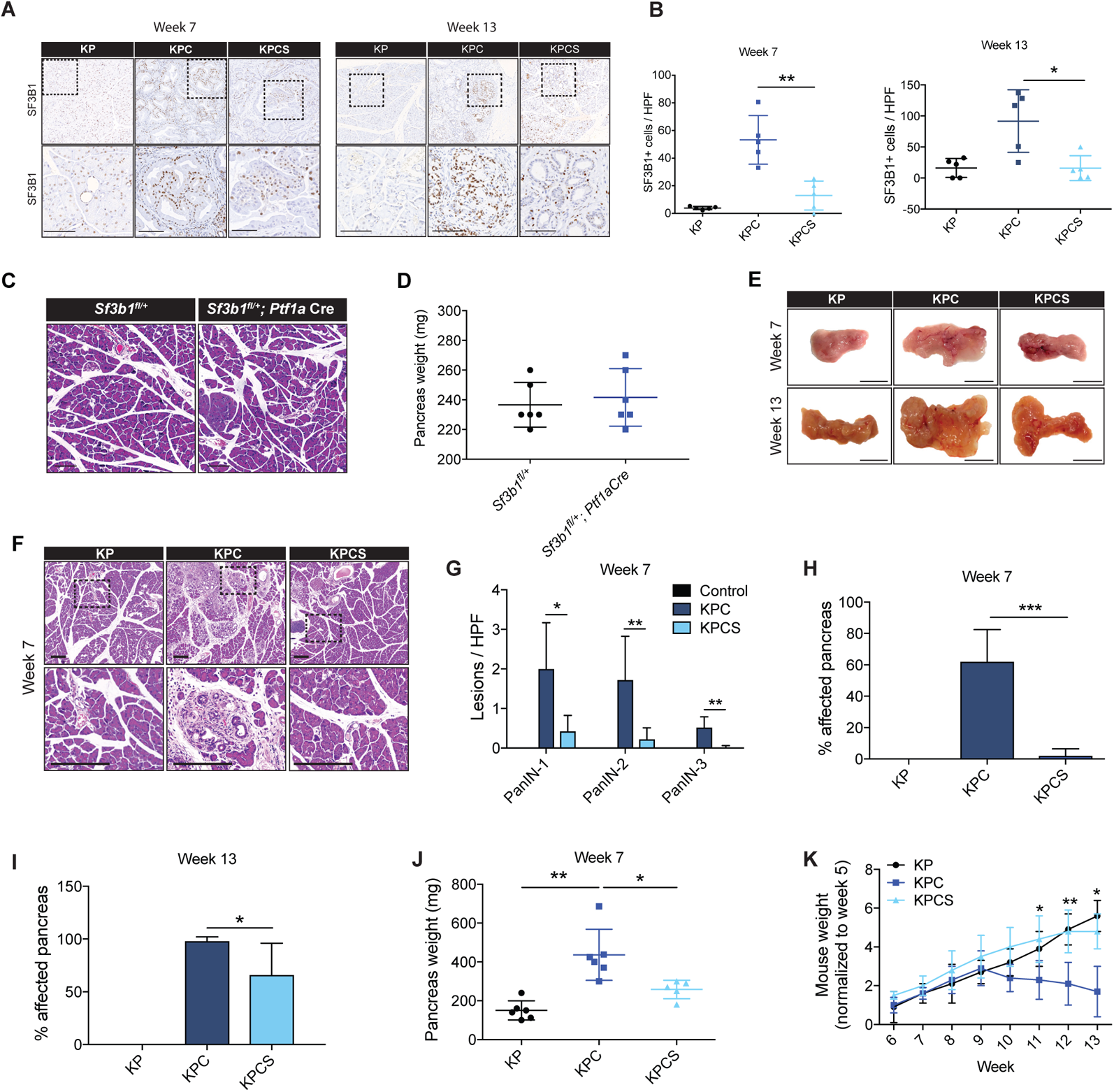
Effect of SF3B1 reduction on PDAC growth. **(A)** Representative immunohistochemical staining for SF3B1 in pancreata from 7- and 13-week-old KP, KPC and KPCS mice. Dashed lines indicate the area magnified below. Scale bar is 100 µm. **(B)** Quantification of SF3B1 positive areas in KP, KPC and KPCS mice at 7 and 13 weeks of age. Ten high power fields (HPF) were analysed per animal (n = 5). * P<0.05. One-way ANOVA followed by a Tukey’s multiple comparison post-test. **(C)** H&E staining of pancreata from *Sf3b1^fl/+^* and *Sf3b1^fl/+^; Ptf1aCre* mice at 13 weeks of age. Representative images, scale bar is 100 µM. **(D)** Evaluation of pancreas weight of *Sf3b1^fl/+^* and *Sf3b1^fl/+^; Ptf1aCre*, mice at 13 weeks of age. The data are represented as mean ± S.D. (n = 6-7). **(E)** Representative photographs of KP, KPC, and KPCS pancreata from 13 weeks-old mice. Scale bar: 1cm. **(F)** Representative images of H&E staining revealing the histopathologic lesions in pancreata of KP, KPC and KPCS mice at the indicated age. Square dashed lines indicate the area magnified in the image below. Scale bar is 100 µm. **(G – I)** Quantification of the prevalence of mPanIN and PDAC in the H&E-stained pancreatic sections of 7-week-old KP, KPC and KPCS mice (G). Quantification of the affected pancreas at the indicated age (H, I). Ten high power fields (HPF) were analysed per animal. The data are represented as mean ± S.D. (n = 5). * P<0.05, ** P<0.01, *** P<0.001. One-way ANOVA followed by a Tukey’s multiple comparison post-test. **(J)** Pancreas weight of KP, KPC and KPCS mice at 7 weeks of age. The data are represented as mean ± S.D. (n = 6-7). * P<0.05, ** P<0.01, *** P<0.001. One-way ANOVA followed by a Tukey’s multiple comparison post-test. **(K)** Body weight of KP, KPC, and KPCS mice at indicated time point over a period of 13 weeks. The data are normalized to body weight of the mice at 5 weeks of age and represented as mean ± S.D. (n = 6-8). * P<0.05, ** P<0.01. One-way ANOVA followed by a Tukey’s multiple comparison post-test.

**Supplementary Figure 5.**
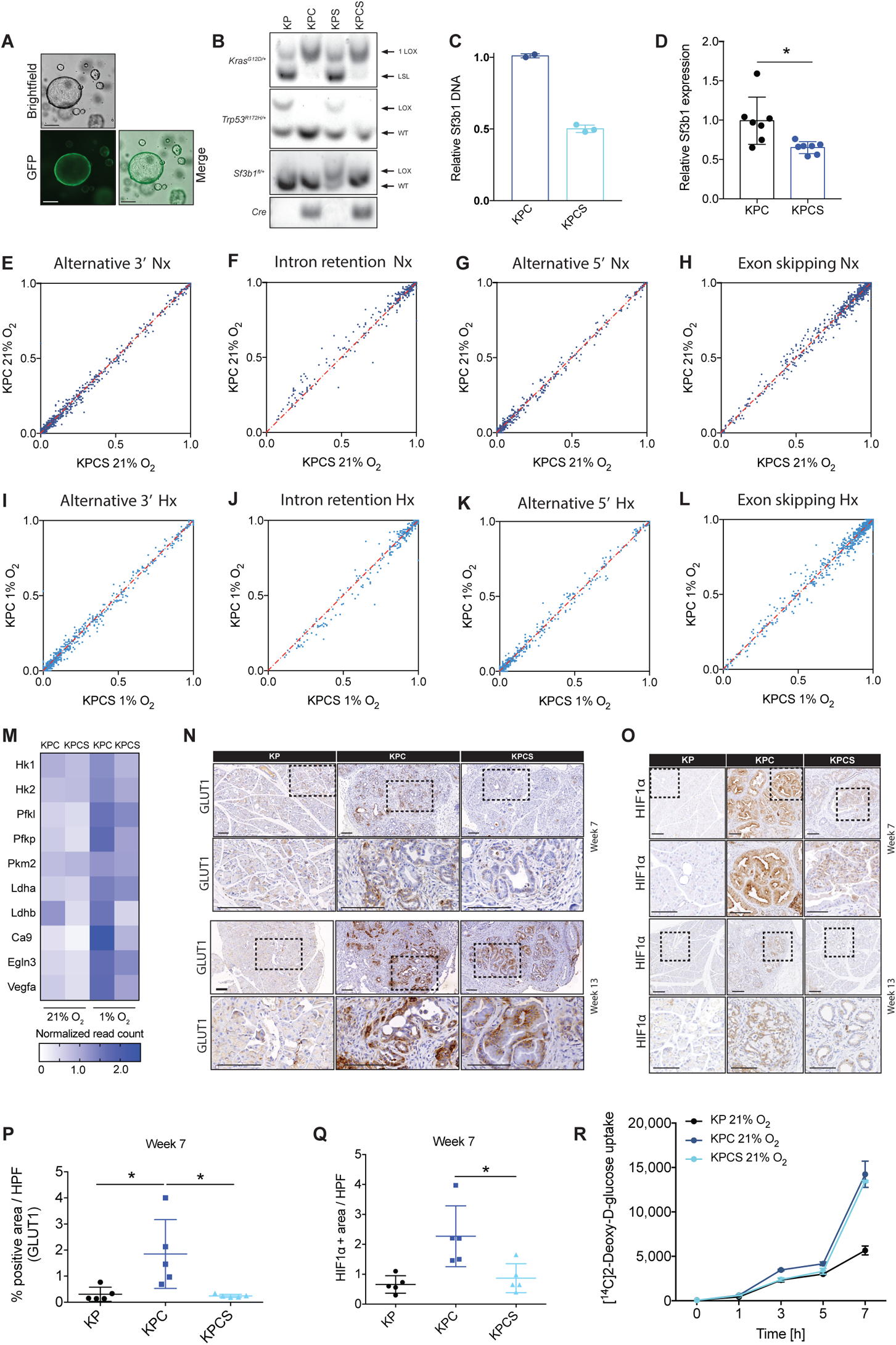
*Sf3b1* heterozygosity affects PDAC growth via HIF signalling. **(A)** Bright field and fluorescence images of mouse organoids transduced with a GFP-Cre virus after FACS. **(B)** Representative PCR analysis of genomic DNA from KP, KPC, KPS and KPCS organoids. The Cre-mediated recombination of the conditional allele *KrasG12D/+*, *Trp53R172H/+* generates a single LoxP site and 2 LoxP sites for *Sf3b1*. (**C**) *Sf3b1* DNA levels at exon 5 for validation of monoallelic Sf3b1 knockout in the indicated organoid lines assessed by qPCR. Values were normalized to *Sf3b1* exon 2 DNA levels, which remain unaffected by Cre-lox recombination. **(D)** *Sf3b1* mRNA expression of KPC and KPCS organoid lines. *P<0.05, ** P<0.01, two-tailed unpaired t-test. **(E-L)** Scatter plot of alternative 3’ end usage (E, I), intron retention (F, J), alternative 5’ end usage (G, K) and exon skipping (H, L) events in KPC and KPCS organoids cultured in normoxia (E to H) and hypoxia (I to L). Events were computed by SplAdder(Kahles et al., 2016), only high confidence events were considered for the analyses. **(M)** Heatmap displaying normalized read counts for the indicated genes derived by RNA-sequencing. **(N)** Representative immunohistochemical staining for GLUT1 in pancreata from 7-week and 13-week old KP, KPC and KPCS mice. Dashed lines indicate the area magnified below. Scale bar: 100 µm. **(O)** Representative immunohistochemical staining for HIF1α in pancreata from 7-week and 13-week old KP, KPC and KPCS mice. Dashed lines indicate the area magnified below. Scale bar: 100 µm. **(P)** Quantification of GLUT1 positive areas in KP, KPC and KPCS mice at 7 weeks of age. Ten high power fields (HPF) were analysed per animal (n = 5). * P<0.05. One-way ANOVA followed by a Tukey’s multiple comparison post-test. **(Q)** Quantification of HIF1α positive areas in KP, KPC and KPCS mice at 7 weeks of age. Ten high power fields (HPF) were analysed per animal (n = 5). * P<0.05. One-way ANOVA followed by a Tukey’s multiple comparison post-test. **(R)** KPC and KPCS organoids were incubated in normoxic conditions in medium containing [^14^C]-2-deoxyglucose at different time-points and processed for glucose uptake measurements. Counts were normalized to the cell number (*n* = 4 biological replicates per time point and condition). ** P<0.01, One-way ANOVA followed by a Tukey’s multiple comparison post-test.

**Supplementary Figure 6.**
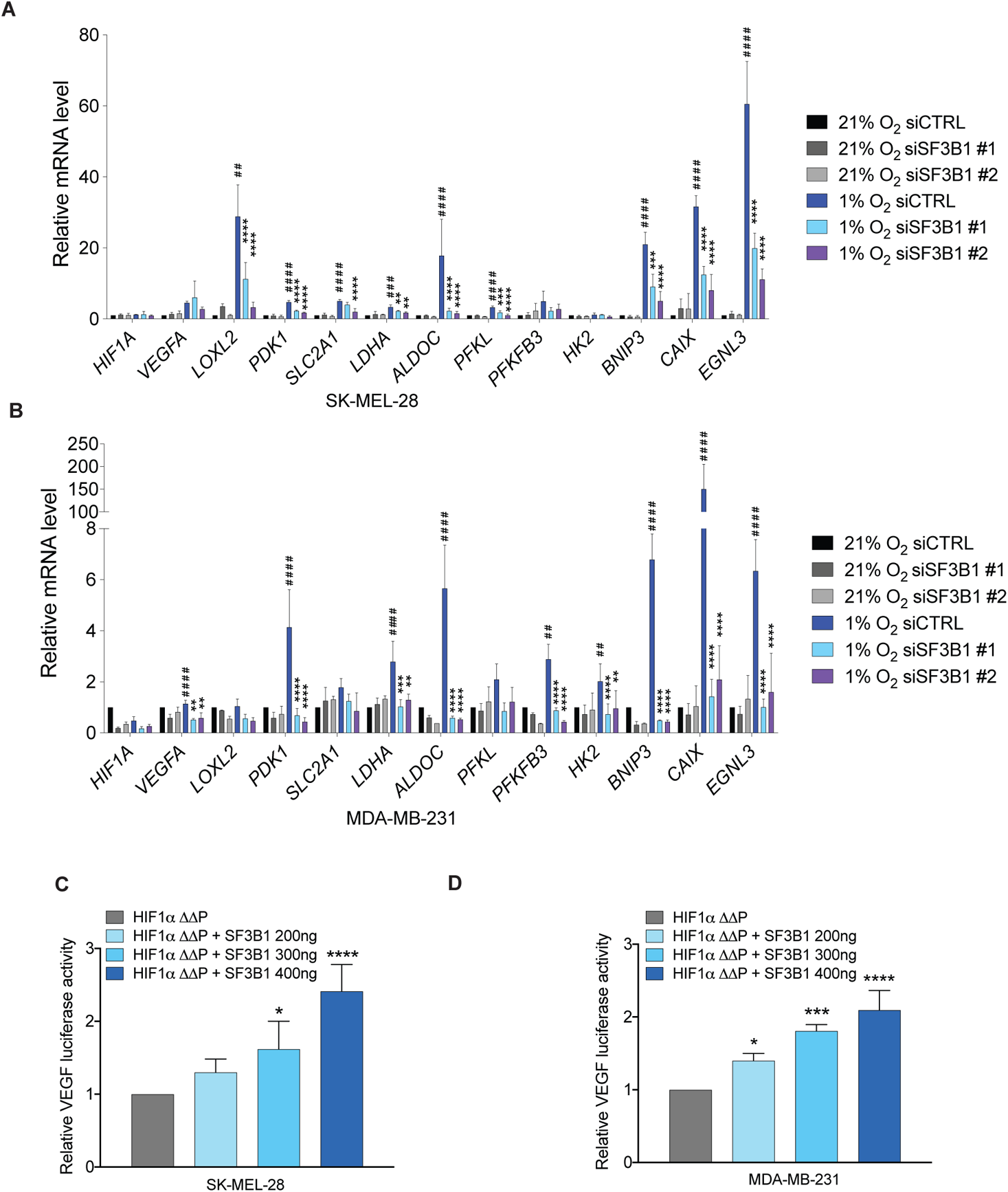
SF3B1 reduction affects HIF signalling in cancer cell lines. **(A and B)** qPCR analyses of the indicated genes in SK-MEL-28 (A) and MDA-MB-231 (B), transfected with the depicted siRNAs and incubated in 21% or 1% O_2_ for 12 hours. Expression levels are relative to *TBP*. Data are shown as mean ± S.D. of biological replicates (n=3). *P<0.05, **P<0.01, ****P<0.0001, data normalized to 1% O_2_ siCTRL. For # P<0.05, ## P<0.01, #### P<0.0001, data are normalized to 21% O_2_ siCTRL. Two-tailed unpaired t-test. **(C and D)** Transfection of a *VEGF* promoter-luciferase reporter alone or together with either HIF1αΔΔP (the constitutively active form of HIF1α), or in combination with different concentrations of SF3B1 plasmid in SK-MEL-28 (C) and MDA-MB-231 cells (D). Data are shown as mean ± S.D. of biological replicates (n = 3). *<P0.05, **P<0.01, ***P<0.001, **** P<0.0001. One-way ANOVA followed by a Dunnet’s multiple comparison post-test.

**Supplementary Figure 7.**
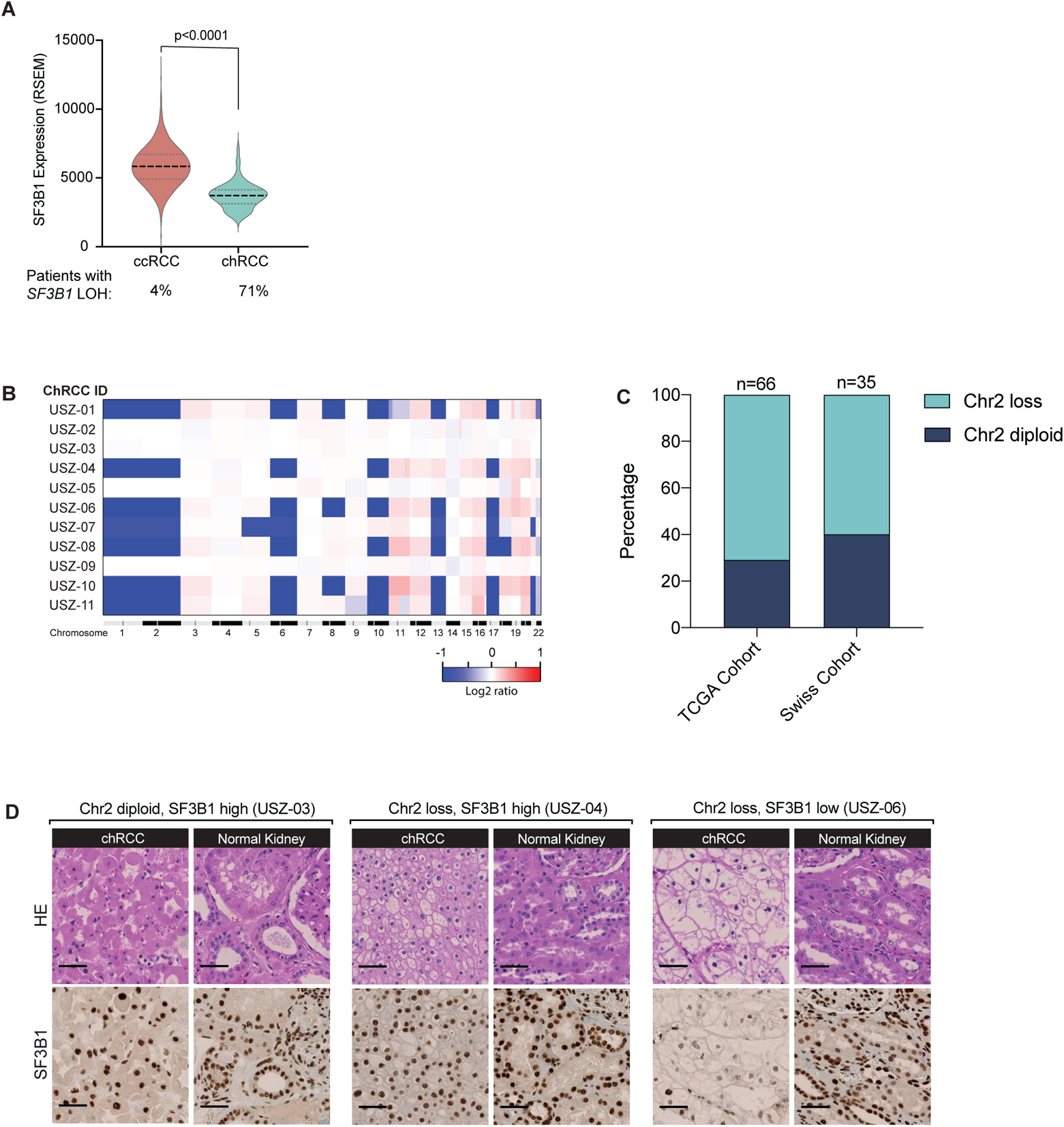
*SF3B1* heterozygosity is frequently observed in chRCC. **(A)** Expression of SF3B1 and loss of heterozygosity (LOH) in clear cell renal cell carcinoma (ccRCC) and chromophobe renal cell carcinoma (chRCC) based on TCGA data. One-way ANOVA followed by a Tukey’s multiple comparison post-test. **(B**) Copy number variations (CNV) analysed with array-based comparative genome hybridization (array-CGH) of tumors from 11 chRCC patients of the Swiss-cohort. **(C)** Chromosome 2 status in chRCC samples of the TCGA cohort and the Swiss cohort. The Swiss cohort is composed of samples published by Ohashi et al., 2019 (n=24) and the samples shown in Suppl. Fig. 5A (n=11). **(D)** Representative H&E and IHC staining of SF3B1 chRCC patients without loss (left) and with loss (middle and right) of chromosome 2. The middle staining shows compensation of heterozygous loss of *SF3B1*, whereas the sample on the right has reduced levels of SF3B1. Scale bar is 50μm.

